# A Multimodal Spatial and Epigenomic Atlas of Human Adult Lung Topography

**DOI:** 10.1101/2025.05.23.655666

**Authors:** Thu Elizabeth Duong, Dinh Diep, Kimberly Y. Conklin, Indy Bui, Jeffrey M. Purkerson, Eric Boone, Jacqueline Olness, Sahil Patel, Beverly Peng, Colin Kern, Zoey Zhao, Ravi S. Misra, Heidie L. Huyck, Jamie M. Verheyden, Zea Borok, Yun Zhang, Richard H. Scheuermann, Quan Zhu, Gail Deutsch, James Hagood, Xin Sun, Kun Zhang, Gloria S. Pryhuber

**Affiliations:** Department of Pediatrics, University of California San Diego, La Jolla, CA; Department of Bioengineering, University of California San Diego, La Jolla, CA; Department of Pediatrics, University of Rochester, Rochester, NY; Department of Cellular and Molecular Medicine, University of California San Diego, La Jolla, CA; Department of Informatics, J. Craig Venter Institute, La Jolla, CA; Division of Pulmonary, Critical Care, Sleep Medicine and Physiology, Department of Medicine, University of California San Diego, La Jolla, CA; National Library of Medicine, National Institutes of Health, Bethesda, MD; Department of Pathology, Seattle Children’s Research Center, Seattle, WA; Department of Pediatrics, Children’s Research Institute, University of North Carolina, Chapel Hill, NC

## Abstract

Developing high-resolution reference maps of disease-susceptible spatial niches is a critical step to mitigating the profound effects of lung disease. Here, we present an integrated multimodal single-nucleus human lung atlas (snHLA) profiling 746,047 nuclei from 49 mapped lung blocks spanning clinically relevant distal airways, alveoli, and interstitium across 11 healthy adults. Integrating snRNA-seq and SNARE-seq2, which co-assays chromatin accessibility and gene expression from the same nucleus, we resolved 70 molecularly distinct populations and captured 332,846 accessible chromatin regions, nominating new transcriptional regulators of human lung cell diversity. Spatial transcriptomics using MERFISH mapped 25 cell populations across 7 structural neighborhoods and multiplexed immunofluorescence localized cell subtypes and distal airway-defining protein markers, expanding and validating distinct lung structure-specific cell populations. This open access snHLA and companion Cell Type and Marker Gene Dictionary with anatomically aligned nomenclature delivers a foundational resource at an unprecedented resolution to interrogate the origins of lung pathophysiology.

## Main

Respiratory disease affects all age groups, communities, and healthcare systems, accounting for 40% of mortality and 10% of disability-adjusted life-years lost globally^1^. Given the ongoing impact of lung disease, it is critical to identify novel diagnostics, predictors of disease severity, therapeutics, and preventative strategies. This requires a high-resolution understanding of the molecular regulation of lung cellular phenotypes within spatial niches where diseases originate. Towards this goal, atlas-building consortia have leveraged single cell transcriptomics to create foundational cellular guides for the healthy adult lung^2–5^. These efforts have advanced our knowledge by identifying novel cell types, subtypes, and states, adding depth to prior knowledge, and substantiating predicted cell types while revealing unexpected variations.

To date, human lung atlases are largely static, one-dimensional single-cell transcriptional snapshots of dissociated intact cells, favoring cell types easily released from surrounding matrix. Profiling of nuclei has been limited. Anatomically, beyond the lung parenchyma, studies have begun to prioritize the trachea, large proximal airways (bronchi; generations 1-5)^3,6–8^, and small distal airways including primary bronchioles (luminal diameter <2 mm; generations 8-11), distal lobular bronchioles (luminal diameter <1 mm; generations 12-16), and terminal and respiratory bronchioles (generation 17 and beyond)^6,8,9^. Emerging spatial single-cell technologies are starting to map molecularly defined cell populations in the airways and parenchyma^3,9,10^.

Our reference adds critical new depth toward building a comprehensive healthy lung cell atlas. Large-scale multimodal profiling of high-quality single nuclei (sn) is presented. Interrogation of nuclei permits prospective study of archived frozen tissues, reduces batch effects, avoids dissociation-induced transcriptional stress, and better represents large, fragile cells such as alveolar epithelial type 1 (AT1) cells^11^. The samples assayed, anatomically mapped and remaining available from the BioRepository for INvestigation of Diseases of the Lung (BRINDL), target airway regions of high clinical interest including the small distal airway transition from bronchi to non-cartilaginous bronchioles. A comprehensive understanding of the distal airways is particularly impactful as many lung diseases, such as cystic fibrosis, asthma, chronic obstructive pulmonary disease (COPD), and idiopathic pulmonary fibrosis (IPF), significantly involve or originate in the distal airways^8,11^. In adults, these small airways are often referred to as “silent” or “quiet” zones as disease starts here without noticeable symptoms or loss of lung function until the condition is advanced and often irreversible^12,13^. Recent microdissection studies of <2 mm bronchiolar airways uncovered unique cellular composition suggesting new targets for interventions^6,8,9^. These studies were limited, however, to cellular transcriptomes and did not capture well the bronchial to bronchiolar junction. Our unified nucleus-centric approach builds on these studies by jointly profiling accessible chromatin (AC), as well as cell marker and transcription factor (TF) gene expression, precisely linking cell molecular identity and gene regulation, focusing on lung regions enriched in the distal airway transition and quiet zones.

Layering complementary ‘omic modalities on sc/snRNA-seq expands previous cell-type-defining insights. TFs are pivotal in cell fate determination and disease pathogenesis, binding to accessible chromatin to activate or repress gene transcription. We applied single-nucleus chromatin accessibility and mRNA expression sequencing (SNARE-seq2)^14^, a co-assay method that employs shared barcodes to accurately align ‘omic profiles. Integrating snRNA-seq and SNARE-seq2 to cross-validate and synergize the strengths of each technology, we profiled 746,047 single nuclei and resolved 70 molecularly distinct cell populations from 49 mapped lung tissue blocks capturing the distal airways (subsegmental bronchi to respiratory bronchioles), alveoli, and interstitium, across 11 healthy, non-tobacco using adult donors. We importantly add spatial context and validation by massively multiplexed error-robust fluorescence in-situ hybridization (MERFISH) and highly multiplexed immunofluorescence (MxIF). In donor matched regions, we mapped 25 cell populations across 7 lung structural neighborhoods using MERFISH, and we validated cell type and airway-level specific molecular markers by MxIF antibody (34-45) panels. Our multimodal single-nucleus human lung atlas (snHLA) will serve as a fundamental resource furthering multi-disciplinary investigations in healthy and pathologic respiratory biology.

## Results

### Building a multi-scale adult snHLA

To systematically capture the cellular organization of the healthy human lung in multiple regions from airways to parenchyma, we developed a streamlined protocol for sc/sn ‘omics to identify and process mapped lung blocks containing representative structures sourced from the accessible and expanding human tissue repository of the NHLBI LungMAP BRINDL (Fig. 1, Methods). We chose 11 transplant-quality lungs from deceased donors aged 19 to 59 years old, spanning average pre-, peak, and post-peak lung function periods, with no reported lung disease or tobacco history. Although strict inclusion criteria were applied to reflect “healthy” adult lung, clinical and pathologic findings are reported for consideration (Supplementary Table 1). By self-report, donors represent 4 females and 7 males, 55% white, 18% black, 18% hispanic, and 9% asian. Based on gross-imaging and systematic re-mapping of sample origin (lung lobe, bronchopulmonary segment, and relative coordinate position), 49 OCT-embedded fresh frozen lung blocks were selected containing target structures confirmed by pathologist (G.D.) review of H&E sections serial to cryosections used for nuclei dissociation (Extended Data Fig. 1a-b, Supplementary Fig. 1 and Table 2). Each block was classified by the largest airway identified and contained all subsequent smaller airways and alveoli. Neighboring tissues, formalin inflated and paraffin embedded (FFPE), were reviewed for structure and pathology (Supplementary Fig. 2). All lung tissues are registered in the Human BioMolecular Atlas Program (HuBMAP) common coordinate framework (CCF) and associated with multi-scale anatomic location (gross to microscopic), detailed clinical metadata, and subjected to standardized processing protocols to enable reproducibility.

**Fig. 1:**
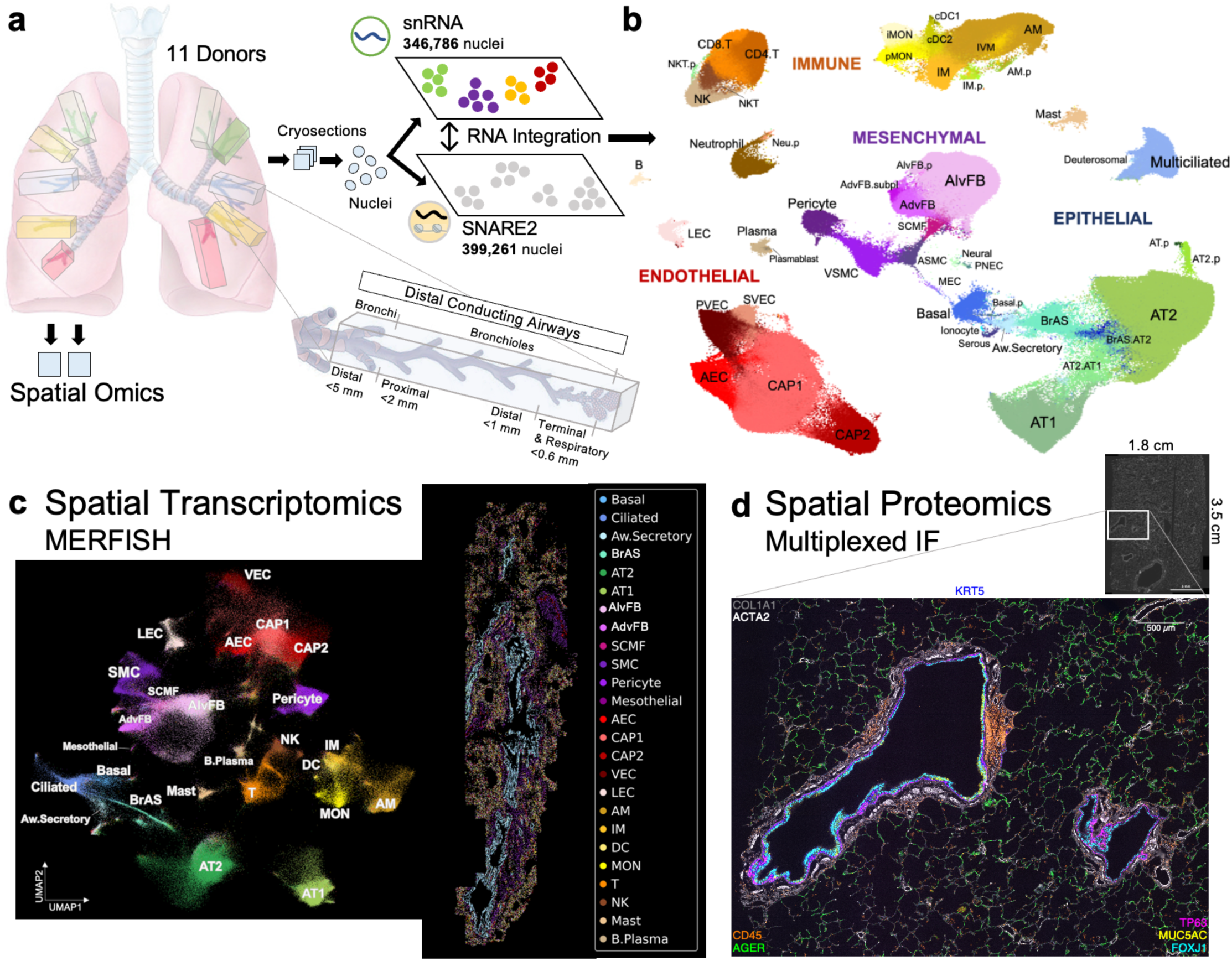
Overview of integrated single-nucleus human lung atlas (snHLA) with spatial multiomic profiling. **a**, Schematic of anatomically mapped, deceased human lung donor blocks processed using 10X snRNA-seq (snRNA), single-nucleus chromatin accessibility and mRNA expression sequencing (SNARE2), multiplexed error-robust fluorescence in situ hybridization (MERFISH), and highly multiplexed immunofluorescence (MxIF). 11 total donors were profiled using one or more technologies. **b**, Combined UMAP embedding of integrated RNA data from snRNA and SNARE2 with aligned cell type annotations (subclass level 3). Also see Supplemental Table 4 for annotation details. **c**, Joint embedding of snRNA and MERFISH identified 25 lung cell types. Left, UMAP projection of 260,272 MERFISH-identified cells. Right, MERFISH spatial map of lung section colored by cell type. **d**, Visualization of eight protein markers in airway and alveolar structures from large lung tissue section using MxIF. Scale bar is 500 µm.

### Foundational single-nucleus grid

To characterize transcriptomic profiles from the distal subsegmental bronchi to terminal and respiratory bronchioles and alveoli, following quality control steps, we identified 346,786 snRNA profiles, consisting of 30% epithelial, 16% mesenchymal, 28% endothelial, 25% immune, and 0.1% peripheral nervous system cells (Fig. 2a, Extended Data Fig. 2, and Supplementary Table 3). We used a stepwise hierarchical approach to annotate populations by cell compartment (class), substructure (subclass.L1), and increasing granularity based on known and sn transcriptome-defined markers (subclass.L2-5) (Methods, Supplementary Table 4). We identified 31 major cell types (subclass.L2) using canonical markers and we further annotated 38 cell subtypes (69 total molecular classifications) based on unique gene signatures (subclass.L5). Aligning with efforts to standardize lung cell nomenclature, we incorporated LungMAP CellCards consensus annotations^15^, Cell Ontology (CL, https://cell-ontology.github.io/), and HuBMAP

**Fig. 2:**
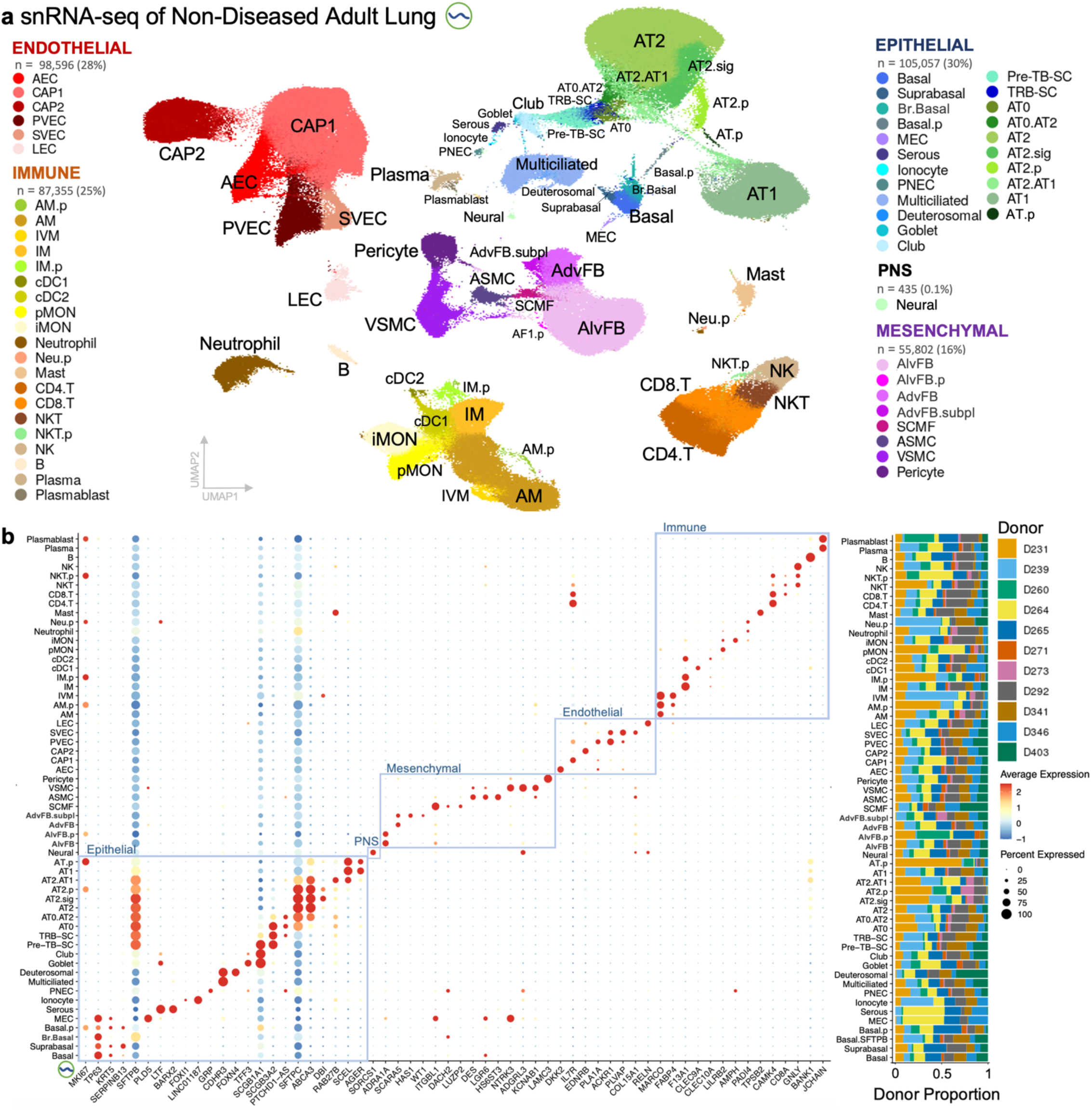
Single-nucleus transcriptomic profiling of human adult lungs captured major cell types along with intermediate and proliferating cell states in the distal airways and lung parenchyma. **a**, UMAP visualization of snRNA-seq dataset consisting of 346,786 nuclei and 58 cell molecular profiles (subclass level 4) in the epithelium, mesenchyme, endothelium, immune and peripheral nervous system. See Supplementary Table 3-5 for QC metadata, description of cell type abbreviations, multilevel annotations with key marker genes, and alignment with Cell Ontology. **b**, Dot plot of known and select marker genes that distinguish annotated cell types (left) and bar plot of proportion of annotated nuclei captured from each donor (right). Gene expression normalized per gene.

Human Reference Atlas language (ASCT+B, Supplementary Table 5)^16,17^. To promote findable, accessible, interoperable, and reusable (FAIR) cell type definitions and to support cell sorting and labeling, we applied NS-Forest, a machine learning method to identify necessary and sufficient marker genes at each annotation level (Methods, Supplementary Tables 4 and 6)^18–20^.

In Fig. 2 subclass.L4 annotations, we captured twelve small airway specific epithelial cell types including submucosal gland (SMG) cells, ionocytes, pulmonary neuroendocrine cells (PNEC), ten distal bronchiolar and alveolar, eight mesenchymal, 6 endothelial, and twenty immune cell types, plus few neural cells. Of these, eight clusters expressed *MKI67, TOP2A*, and *CIT*, consistent with a proliferative state. In alignment with snRNA markers, spatial proteomics using 34 to 45 protein MxIF antibody panels mapped diverse cell types and structures in donor matched lung sections (Fig. 3, Extended Data Fig. 3-4, Supplementary Fig. 3 and Table 12). In the epithelium, *TP63*^+^ cells subclustered into *KRT5*^+^ basal, *KRT5^+^SERPINB13^+^*suprabasal, *KRT5^+^MKI67+* proliferating basal (basal.p), *KRT5^+^ACTA2^+^PLD5^+^*SMG myoepithelial cell (MEC), and *SFTPB^+^* distal bronchiolar basal cells (br.basal). In addition to SMG MECs, we identified SMG *LTF^+^BARX2^+^PRB4^+^* serous cells. We detected *DEUP1^+^FOXN4^+^* deuterosomal cells, described as precursors to *FOXJ1^+^CDHR3^+^RSPH1^+^*multiciliated cells^21^. At subclass.L2, *SCGB1A1* and *SCGB3A2* expression split into *SCGB1A1^+^SCGB3A2^-^* and *SCGB3A2^+^SFTPB^+^*populations classified as conducting airway secretory (aw.secretory) and bronchiolar airway secretory (BrAS) cells, respectively. Iterative sub-clustering of all epithelial airway cells divided *SCGB1A1^+^SCGB3A2^-^*aw.secretory cells into *TFF3^+^MUC5B^+^MSMB^+^*goblet and *ERN2^+^WNK2^+^* club cells. *SCGB3A2^+^SFTPB^+^*cells divided into *SCGB1A1^+^* pre-terminal bronchiole (pre-TB-SC), *SCGB1A1^-^* terminal and respiratory bronchiole secretory cells (TRB-SC), *SCGB1A1^-^SFTPC^+^*bronchiolar alveolar type 0 (AT0), and *SCGB1A1^-^SFTPC^+^ABCA3^+^*intermediate AT0 - alveolar epithelial type 2 (AT0.AT2) cells^9^. In the alveoli, we classified *ABCA3^+^* AT2 cells, including *ABCA3^+^FABP5^+^DBI^+^*AT2 signaling cells (AT2.sig)^5^ and *ABCA3^+^SCEL^+^RAB27B^+^*intermediate AT2.AT1 cells, *AGER^+^SCEL^+^* AT1 cells, and *MKI67^+^TOP2A^+^CIT^+^* proliferating *ABCA3^+^* AT2 (AT2.p) and *ABCA3^+^AGER^+^* alveolar epithelial transitional (AT.p) populations.

**Fig. 3:**
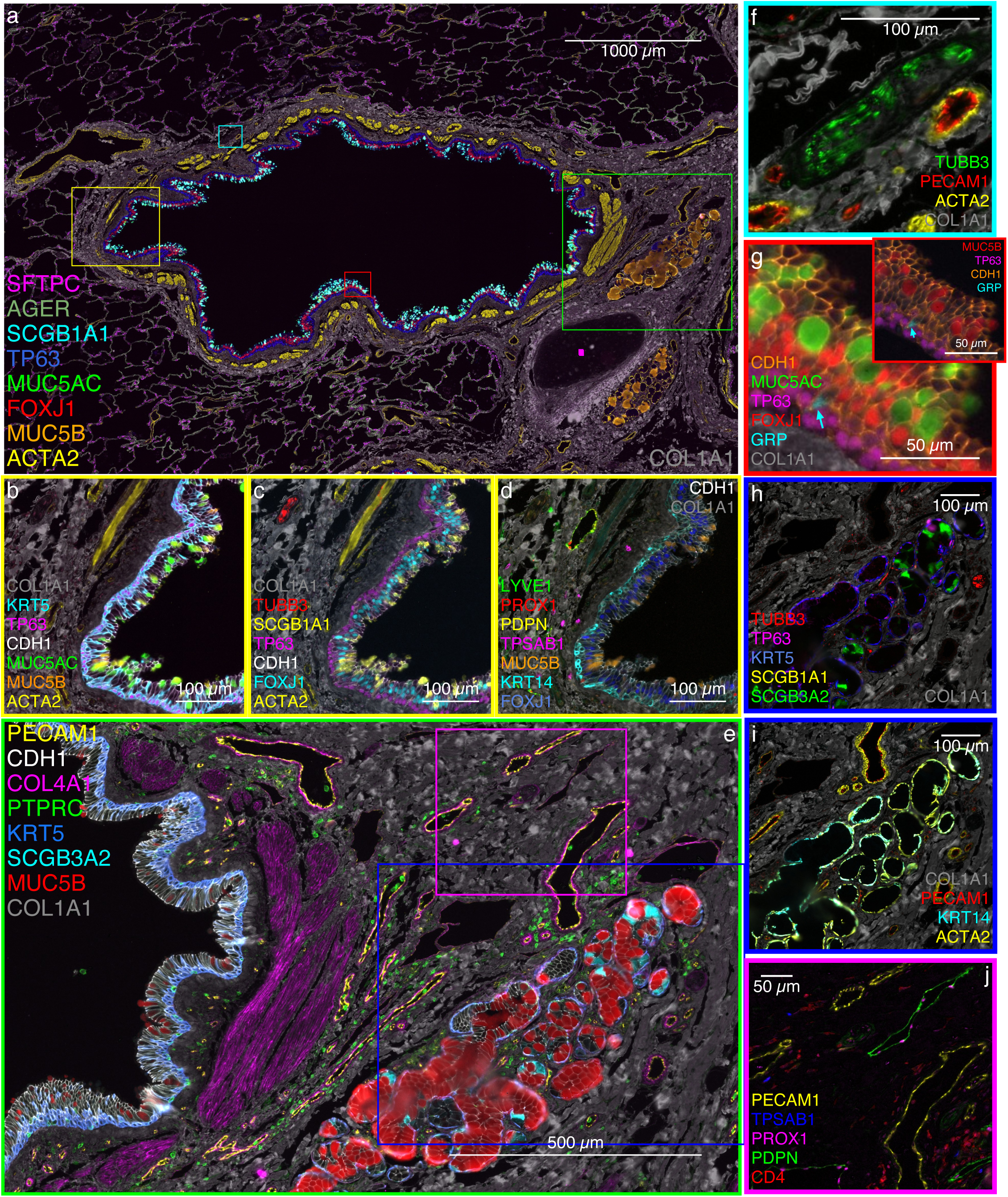
Spatial proteomics resolves structural and cellular diversity in human adult lung. Representative lung cell types identified by parallel single nuclei transcriptomics (snRNA-seq and SNARE-seq2) and localized by antibody-mediated, protein marker-based multiplexed immunofluorescence (MxIF). **a**, Distal bronchus (I.D. 1-2 mm) with cartilaginous plate (pink diamond) and regions of interest (ROI yellow panels **b-d**, green panel **e**, turquoise panel **f**, red panel **g**). **b**, Demonstrates cuboidal basal cells with TP63*^+^* nuclei (pink) and KRT5*^+^*cytoplasm (turquoise); also more columnar KRT5^+^ cells extend toward the airway surface; CDH1 marks apical airway epithelial cell junctions (white); mucin (MUC5AC, green and fewer MUC5B, orange) secreting goblet cells; airway and vascular smooth muscle (ACTA2, yellow); and collagen-rich (COL1A1, gray) extracellular matrix (ECM). **c**, Demonstrates TP63^+^ basal cells (pink); transcription factor FOXJ1^+^ multi-ciliated cell nuclei (turquoise); many of the mucin positive cells in panel **b** are also SCGB1A1^+^ (yellow); TUBB3^+^ nerve cell bodies (red) with nuclei and fibers are also detected. **d**, Lymphatic endothelial cell membrane (PDPN, yellow) and nuclei (PROX1, red); LYVE1 (green) is also noted in lymphatic endothelial cells and in interstitial cells that are CD163^+^ (not shown). Basal cells are heterogeneously KRT14^+^ (turquoise); MUC5B^+^ (orange) goblet cells are more clearly shown; TPSAB1^+^ (pink) mast cells are scattered in the ECM. **e**, Bronchial airway and submucosa extracted from panel **a**; membrane protein CDH1 (white) segments the pseudo-columnar airway epithelium; collagen 4A1 (COL4A1, magenta) is selectively identified forming basement membrane for the smooth muscle bundle of the muscularis mucosa (ACTA2^+^ in panel **a**), surrounding the cross-sectioned small vessel PECAM1^+^ endothelium in the lamina propria just below the epithelium, and in longitudinal blood vessels external to the smooth muscle and in the adventitia to the right of the SMG; submucosal gland (SMG) secretory cells vary in their content; panel **e** demonstrates MUC5B (red) predominance with scattered cells containing SCGB3A2 (turquoise); scattered MUC5B cells also occur in the airway epithelium to the left. **f**, Demonstrates a TUBB3^+^ nerve bundle (green) embedded in COL1A1^+^ ECM and in the neighborhood of muscularized vessels. **g**, Digital zoom to demonstrate single pulmonary neuroendocrine cell (GRP, turquoise arrow) in proximity of basal (TP63, pink) and multiciliated cells (nuclear FOXJ1, red); additionally, goblet cells containing both MUC5AC (green) and MUC5B (inset, red) are demonstrated. **h**, Identifies cytoplasmic SCGB3A2^+^ (green) cells in SMG epithelium but not airway epithelium; SCGB1A1 (yellow) in airway epithelium (see panel **a**) but not SMG; nerve bundles and fibers (TUBB3, red) are adjacent and within the SMG; **h-i**, SMG myoepithelial cells are consistently TP63^+^ (pink), KRT5^+^ (blue), KRT14^+^ (turquoise) and ACTA2^+^ (yellow), as compared to airway basal cells that are intermittently KRT14^+^ and ACTA2 negative (see panel **d**). **j**, This inset highlights ECM-embedded lymphatic endothelium (nuclear PROX1, pink and membrane PDPN, green) as compared to PECAM1^+^ (yellow) vascular endothelium; scattered submucosal mast cells (TPSAB1, blue) and loosely clustered CD4^+^ T cells (red) occuring near vasculature. Image based on D273.

Beyond the epithelium, we detected *ADRA1A^+^GRIA1^+^PCDH15^+^*alveolar fibroblasts (AlvFB) and *SCARA5^+^MFAP5^+^* adventitial fibroblasts (AdvFB) along with an AdvFB subpopulation *HAS1^+^WT1^+^*subpleural fibroblasts (AdvFB.subpl). *HAS1^+^* fibroblasts localized to subpleural regions^22^. Besides WT1 expression, AdvFB.subpl lacked additional mesothelial cell markers such as *UPK3B, FREM2*, and *CALB2*. In alignment with CellRef^2^, we identified *ITGBL1^+^PDGFRA^+^FGF18^+^DACH2^+^*secondary crest myofibroblasts (SCMF). These lack *ASPN, CDH4, HHIP, LGR5, LGR6* used to identify alveolar ductal myofibroblasts and peribronchial fibroblasts in mouse^23–25^. Endothelial cells included *DKK2^+^GJA5^+^*arterial endothelial cells (AEC), *FCN3^+^IL7R^+^* capillary general endothelial cells (CAP1), *HPGD^+^EDNRB^+^* capillary aerocyte endothelial cells (CAP2), and *ACKR1^+^* venous endothelial cells (VEC) which separated into *HDAC9^+^PLAT^+^PLA1A^+^*pulmonary venous and *COL15A1^+^PLVAP^+^ABCB1^+^*systemic venous endothelial cells (PVEC, SVEC, respectively)

Monocytes and macrophages expressed *CD163*. *BACH1^+^MRC1^+^MSR1^+^*macrophages consisted of *PPARG^+^MARCO^+^INHBA^+^*alveolar macrophages (AM), *MKI67^+^* proliferating AM (AM.p), *FABP4^+^* intravascular macrophages (IVM)^26^, *F13A1^+^*interstitial macrophages (IM), and *MKI67^+^* proliferating IM (IM.p). *FCN1^+^* monocytes divided into *LILRB2^+^FCGR3A^+^*non-classical or patrolling monocytes (pMON) and *VCAN^+^AMPH^+^*classical or inflammatory monocytes (iMON). Additional myeloid cells consisted of *CSF3R^+^IL1R2^+^GLT1D1^+^PADI4^+^*neutrophils, *MKI67^+^* proliferating neutrophils (Neu.p), and *TPSB2^+^KIT^+^MS4A2^+^* mast cells. Immune lymphoid cells comprised of *CAMK4^+^* T cells, *CD3D^-^ GNLY^+^KLRD1^+^NCAM1^+^* natural killer (NK) cells, *BANK1^+^* B cells, *JCHAIN^+^* plasma, and *JCHAIN^+^MKI67^+^*plasmablasts. *CAMK4^+^* T cells divided into *CD4^+^*and *CD8A^+^CD8B^+^* T cells (CD4.T and CD8.T) and *CD3^+^GNLY^+^KLRD1^+^NCAM1^+^*natural killer T cells (NKT) including *MKI67^+^* NKT.p cells.

In summary, our anatomically targeted, multilevel annotation approach produced a nucleus-centric adult human lung cell and marker gene dictionary for 58 major lung cell types and subtypes at subclass.L4 from disease-relevant small airways and alveoli. Our snRNA grid adds cellular and transcriptomic depth to key anatomic regions during homeostasis identifying small bronchiolar-specific basal and BrAS cell subsets without artifact of microdissection and contributes new profiles for intermediate AT0.AT2, subpleural adventitial fibroblasts, neural cells, and proliferating states (basal.p, AT.p, AF1.p, AM.p, IM.p, NKT.p, neu.p, and plasmablasts).

### Active transcription factor regulators of lung cell heterogeneity

Applying our systematically annotated snRNA grid, we precisely aligned SNARE2 molecular cell types and corresponding regulatory profiles (Methods). Because pivotal cell-type specific regulatory landscapes in human lung are lacking, we utilized SNARE-seq2 to co-assay mRNA and AC from the same nucleus. We profiled 27 lung blocks from seven adult donors (Extended Data Fig. 1a and Supplementary Table 2). Nuclear suspensions from 22 blocks were split to generate matched snRNA and SNARE2 datasets. Over 318,000 SNARE2 nuclei passed both RNA and AC QC thresholds (Methods). Joint SNARE2 RNA-AC profiles were obtained for all annotated clusters in subclass.L3 and subclass.L4 except for rare cDC1s (48/49 and 55/56 clusters, respectively; Fig. 4a subclass.L3, Supplementary Table 8). In addition to cDC1 cells, SNARE2 subclass.L5 did not include arterial endothelial subset 2 cells (AEC-2) but integrated transcriptomic clustering identified intermediate multiciliated and airway secretory cells (multicil.aw.sec) which were not detected in snRNA alone. For downstream analysis, we performed enhanced peak calling using subclass.L3-5 integrated transcriptomic annotations and modified k500 infomap clusters (Methods). We detected 332,846 accessible regions. For subclass.L3, clusters with less than 10 nuclei were removed from differential accessible regions (DAR) analysis (NKT.p and plasmablasts). Fig. 4b depicts unique DAR profiles for the remaining 47 cell types. As related TFs bind to similar transcription factor binding sites (TFBS), we refined predicted cell-type specific TFs by identifying enriched TFBS with corresponding TF gene expression in at least 10% of nuclei. We refer to these transcribed TFs existing in nuclei with corresponding TFBS in open chromatin as active TFs. Dot plots of TF activity in Fig. 4c present top differentially active TFs in subclass.L3 epithelial, mesenchymal, endothelial, and immune cells. Supplemental Tables 14-16 provide DARs and TF predictions for differentially enriched TFs and active TFs.

**Fig. 4:**
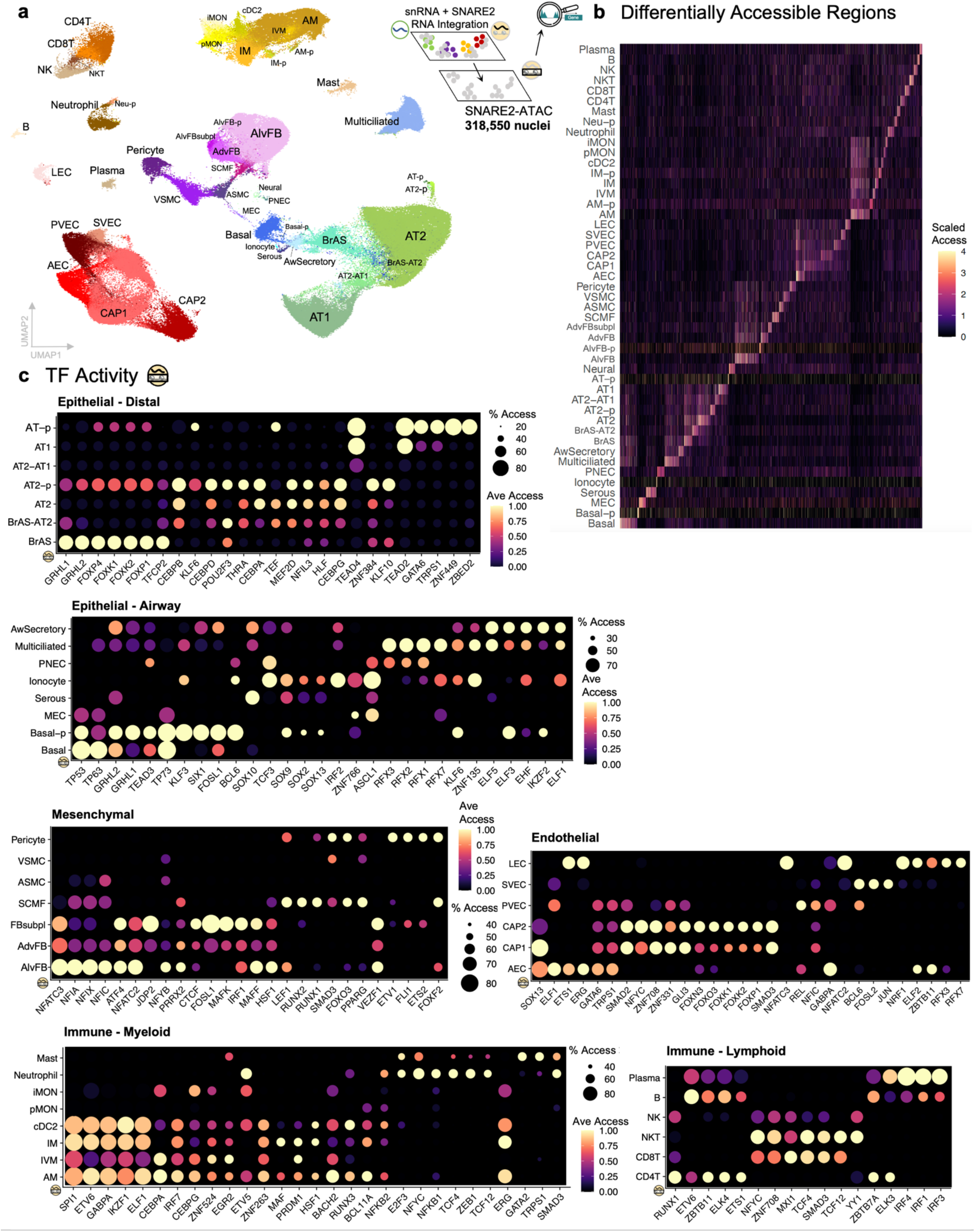
Regulation of human lung cell heterogeneity. **a**, UMAP visualization of >300,000 SNARE2 nuclei with simultaneously profiled RNA and ATAC passing quality control thresholds. Cell type labels (subclass level 3) were generated from integration of snRNA-seq and SNARE-seq2 transcriptomic profiles. See Supplementary Table 4 for cell type definitions. **b**, Heatmap of scaled accessibility for differentially accessible regions for each cell type. **c**, Dot plots of chromVAR average transcription factor (TF) motif accessibility and proportion of cells accessible for top differentially active TFs between cell types in different lung cellular compartments with expression detected in at least 10% of nuclei. See Supplementary Table 9 for cell-type specific differentially accessible regions and ChromVAR TF enrichment.

In the alveolar epithelium, consistent with previous studies, CEBP and TEAD family TFs were most active in AT2 and AT1 cells respectively^27–30^. Proliferating alveolar epithelial cells demonstrated an expanded TF regulatory program compared to their more quiescent counterparts. AT2.p were notably enriched for THRA activity which has been shown to regulate AT2 cell proliferation in mice after injury^31^, and for KLF10 activity implicated in modulation of lung fibrotic processes^32^. In addition to TEAD, zinc finger TFs (ZNF449, ZBED2, and GATA6) and KLF6 were active in AT.p cells. Intermediate AT2.AT1 cells had minimal to no overlapping TF activity with AT2 cells and decreased TEAD4 activity compared to AT1 cells. BrAS cells were enriched in grainyhead (GRHL1, GRHL2, TFCP2), which activate NOTCH1 and NKX2-1 transcription, and forkhead box (FOXK and FOXP) family TFs, which regulate Wnt/β-catenin-dependent transcription. NOTCH and WNT signaling have been demonstrated to regulate *SCGB3A2^+^*respiratory airway secretory (RAS) cells to AT2 cell differentiation^6^. In the airway epithelium, as expected, basal cells were enriched in TP63 activity. In SMG cells, MECs were enriched for ASCL1, a canonical marker of rare PNECs which have also been identified in human SMGs to signal nearby MECs^33^. SOX10 was active in SMG serous cells, consistent with protein staining^34^. More recently identified pulmonary ionocytes were enriched in TCF3, IRF2, and ASCL1. TCF3 has been shown to regulate ASCL1 activity by preventing ASCL1 degradation^35^. Multiciliated cells were enriched in regulatory factor X (RFX) TFs which promote airway ciliogenesis and form a transcriptional complex with FOXJ1^36^. Aw.Secretory cells were active in E74-like factors (ELF), particularly ELF3 and ELF5 which regulate lung epithelial cell development and differentiation^37^.

In the mesenchyme and endothelium, AlvFB, AdvFB, and AdvFB.subpl demonstrated similar enrichment programs including Nuclear Factor of Activated T cells (NFAT) and Nuclear Factor I (NFI) TFs which regulate lung fibroblast proliferation, differentiation, and extracellular matrix (ECM) production^38^. Activating protein-1 (AP-1) TFs (FOXL1, JDP2) involved in selecting enhancer elements to direct fibroblast specialization were enriched in AdvFB.subpl^39^. SCMFs were active in runt-related TFs (RUNX1, RUNX2), which mediate fibroblast-to-myofibroblast differentiation^40^, and in LEF1, a downstream effector of WNT signaling activated during SCMF differentiation^41^. ETS family and subfamily TFs were enriched in pericytes (ETV1, FLI1, ETS2) and AECs (ELF1, ETS1, and ERG). FLI1 modulates pericyte inflammatory responses^42^. ERG1 regulates endothelial cell-cell communication^43^. Flow-sensitive SOX13 was active in AEC and CAP1. Compared to CAP1, CAP2 contained more active FOX TF binding sites (FOXN3, FOXO3, FOXK1, FOXK2, FOXP1). In venous endothelial cells, shear stress activated TFs such as REL were enriched in PVEC and AP-1 TFs (FOSL2, JUN) in SVEC^44^. LECs were active in ETS1, ERG, NFATC3, NFATC2. ETS and NFAT family TFs have been linked to lymphatic identity and function^45^.

In the immune compartment, TF activity profiles largely overlapped for myeloid macrophage and dendritic cell populations with enrichment for SPI, GABPA, IKZF1, and ELF1 binding sites. Continuous SPI1 expression has been shown to favor macrophage over granulocyte differentiation^46^. SPI1 is also required for conventional DC formation and function^47^. AM and IVM were enriched for CEBPA, required for macrophage activation^48^. IM contained high MAF activity, a TF implicated in lung IM development^49^. Neutrophils demonstrated enrichment in TCF4 and TCF12, T-cell factor TFs, NYFC, NFKB, and ZEB1, that regulate Wnt signaling and neutrophil differentiation^50^. Mast cells were enriched for GATA family TFs GATA2 and TRPS1, which have been shown to maintain mast cell identity and cytosolic calcium concentrations fundamental to mast cell function^51^. Notable top active TFs for CD4.T cells included RUNX1, which regulates the survival and homeostasis of CD4+ T cells^52^, and ZBTB7A, which acts redundantly with ZBTB7B to maintain CD4.T cell integrity and effector responses^53^. CD8.T, NKT, and NK are cytotoxic cells and shared signals for NFYC, ZNF708, and MXI1. YY1 was most active in NKT cells, ETV6 in B cells, and IRF4 in plasma cells, which have all been shown to regulate genes essential to corresponding cell type function^54–56^. In summary, our multimodal approach generated tiered resources that validates known TFs and proposes new regulatory candidates for diverse human lung cell types.

### Spatial mapping of lung cell neighborhoods and cell-cell communication

As the multiomic cellular composition of the human lung becomes more granular, adding spatial context to understand how cells communicate and cooperatively function within lung structures is a vital next step. To spatially resolve cellular, molecular, and proteomic heterogeneity in lung structural neighborhoods, we applied imaging-based fluorescence in situ hybridization using MERFISH (Vizgen MERSCOPE) to measure expression of 503 genes, complemented by MxIF staining using the PhenoCycler-Fusion v.2 system (Akoya Biosciences) to visualize up to 45 proteins in lung tissue sections from 6 donors (Fig. 1c-d, Methods, Supplementary Table 2). For MERFISH, we designed and applied a panel of probes to target cell lineage, cell type and subpopulation, TF, signaling, and ECM genes (Supplementary Table 10). After nuclei segmentation and adaptive filtering, we obtained 25.2 million transcripts from 260,272 cells across 2 experiments (Replicate 1: 68,600 cells, 54 transcripts/cell; Replicate 2: 191,672 cells, 70 transcripts/cell). MERFISH RNA transcripts demonstrated high correlation with lung bulk RNA-seq reference data and between replicates (Extended Data Fig. 5a-b). Next, using Harmony, we integrated MERFISH and snRNA to predict MERFISH cell types using modified subclass.L3 labels (Methods). Spatial plots of genes and predicted cell types, correlation with snRNA-seq, and lung marker gene analysis mapped 24 distinct MERFISH lung cell populations (Fig. 1c, Extended Data Fig. 5c-d). The percent of cell types detected in MERFISH was comparable to snRNA-seq (Extended Data Fig. 5e) with the addition of WT1+UPK3B+GAS1+ mesothelial cells, likely due to differences in tissue composition and ability to use both spatial and gene expression to localize these cells to the pleural surface. Localization of key genes for additional validation produced expected spatial patterns (Extended Data Fig. 5f).

### MERFISH recapitulates lung structural neighborhoods

As a proof of concept, we tested whether MERFISH could deconvolve lung structures and expected resident cell types. We applied BANKSY (Building Aggregates with a Neighborhood Kernel and Spatial Yardstick), an algorithm that uniquely combines cell typing and tissue domain segmentation^57^. A clustering resolution was chosen based on the goal of identifying bronchial vs bronchiolar airways (Methods, Supplementary Fig. 4). At this resolution, we categorized 7 lung structural neighborhoods (Fig. 5a, Extended Data Fig. 6). The resulting Bronchiolar Epithelium neighborhood clustering contained a higher percentage of BrAS cells compared to Bronchial Epithelium with many basal, aw.secretory, and multiciliated cells. The majority of AT1, AT2, CAP1 and CAP2 were captured in the Alveolar Neighborhood. SMC, VEC, LEC, and immune cells including B.Plasma contributed to Airway Adventitia. Vascular Bundle neighborhoods contained SMCs and AECs. Mesothelial cells were largely restricted to the Pleura. Overall, cell type composition in lung spatial neighborhoods were as expected, except for AdvFBs, which were enriched in the Interlobular Septa but also present in Airway Adventitia and, to a lesser extent, Alveolar regions.

**Fig. 5:**
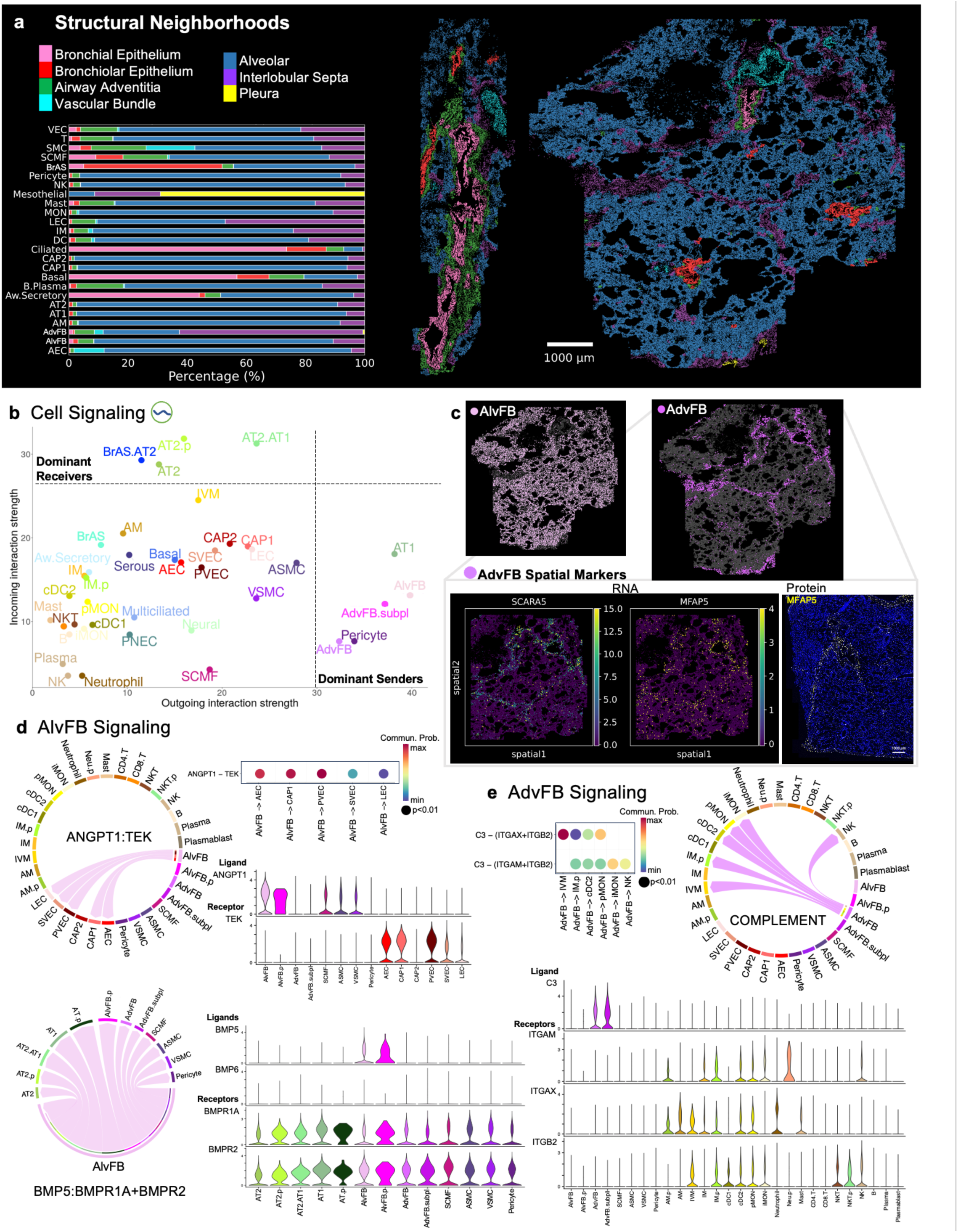
Lung structural neighborhoods and cell communication highlight fibroblast differences. **a**, Bar plot of percentage of MERFISH-identified cell types assigned to each lung structural neighborhood. MERFISH spatial maps of two lung sections colored by neighborhood. **b**, Scatter plot of incoming and outgoing interaction strength highlights dominant senders (sources) and receivers (targets) at subclass level 3. **c**, Spatial map of alveolar fibroblasts (AlvFB) and adventitial fibroblasts (AdvFB) in MERFISH lung section (top). Spatial maps of MERFISH AdvFB-specific marker genes and multiplexed IF AdvFB-specific protein. Chord diagrams and dot plots illustrate significant signaling pathways with communication probabilities of specific ligand-receptor pairs and corresponding cell-cell interactions for AlvFB-directed ANGPT and BMP pathway signaling (**d**) and AdvFB- directed Complement pathway signaling (**e**). Violin plots quantify snRNA expression of ligands and receptors.

### Global lung cellular communication patterns

To further dissect how the complex lung cellular and structural architecture orchestrates physiologic function, we applied CellChat to our snRNA reference and characterized global and local cell-cell interactions. Ligand-receptor interactions are important to homeostasis and dysregulated in disease. These interactions also represent potential therapeutic targets. To gain a high-level perspective of distal lung signaling, we used CellChat to infer communication networks (Methods)^58^. At subclass.L3, CellChat recognized four coordinated signaling patterns each for outgoing and incoming communication (Extended Data Fig. 7a). Cell types clustered into signaling patterns largely by lineage. Next, we defined dominant senders and receivers based on weighted communication probabilities (Fig. 5b). AT1, AlvFB, AdvFB, AdvFB.subpl, and pericytes were major signaling sources. AT2, AT2.p, BrAS.AT2, AT2.AT1 cells were dominant receivers. Heatmaps in Extended Data Fig. 7b list all significant pathways that contributed to outgoing and incoming signals for each group (also Supplementary Table 11).

### Spatial and signaling divergence of fibroblasts

As fibroblasts serve as integral signaling centers yet remain poorly defined spatially and molecularly, we further compared alveolar and adventitial fibroblasts. The majority of AlvFB cells were present in the Alveolar Neighborhood, consistent with previous descriptions (Fig. 5a)^2,5^. In CellRef, alveolar fibroblast 1 cells (AF1), largely curated from single-cell studies, are defined by *TCF21, WNT2*, and *PCDH15*^2^. Our AF1 cells aligned with CellRef AF1s (Extended Data Fig. 2b), but in addition to *PCDH15*, distinguishing sn genes were *ADRA1A, GRIA1*, and *ITGA8*, with low but positive expression of *TCF21* and *WNT2*.

Compared to AlvFB cells, the spatial and molecular characterization of AdvFB cells is less clear. Human *SCARA5^+^MFAP5^+^*fibroblasts have been described in the alveolar mesenchyme (alveolar fibroblast 2, AF2) or adventitia (vascular adventitia and nearby airways – adventitial fibroblast)^4,5,15^. Neighborhood analysis localized *SCARA5^+^MFAP5^+^* fibroblasts to the Interlobular Septa (∼60%), Alveolar (∼20%), Airway Adventitia (∼10%), and Vascular Bundle (∼4%) (Fig. 5a), consistent with MERFISH RNA spatial maps and MFAP5 antibody staining (Fig. 5c), revealing an expanded spatial pattern with a large presence in interlobular septa extending to the pleura. Vascular and lymphatic endothelial cells are also seen in the region of MFAP5+ cells in the interlobular septa. Based on Cell Ontology definitions, adventitia refers to an outermost connective tissue covering of an organ, vessel or other structure. Applying this definition to the lung anatomically aligns with AdvFB found in connective tissue surrounding the airways, vasculature (arteries, veins, and potentially capillaries in alveoli), and lymphatics.

As dominant signaling sources, AlvFB and AdvFB shared outgoing programs involving ECM (collagen, laminin, fibronectin), cell-cell adhesion (*PTPRM*), and neuronal growth factor (*NEGR*) signaling (Extended Data Fig. 8). We highlighted unique pathways for AlvFB and AdvFB with therapeutic potential in lung disease (Fig 5d-e). AlvFBs served as the major source of angiopoietin-1 (*ANGPT1*) signaling to tyrosine kinase receptor (*TEK*) expressed by all endothelial cells except for CAP2. *ANGPT1* is required to maintain endothelial stability and can provide vascular protection, including from acute lung injury^59–61^. AlvFB and AlvFB.p cells expressed bone morphogenic protein 5 (*BMP5*) with high signal communication probability to alveolar epithelial cells and connective tissue cells via *BMPR1A* and *BMPR2*. AdvFB cells were predicted to signal intravascular macrophages (IVM) and other immune cells via *C3* and the complement pathway. AdvFB signaling probability to epithelial BrAS, BrAS.AT2, and AT2.AT1 cells was high via the tenascin family of ECM proteins, implicated in maintaining lung structure and function and providing an advantage in lung aging^62^. By integrating our snRNA reference with spatial transcriptomics and in-silico ligand-receptor tools, we generated a comprehensive, multicellular communication map of the distal lung during homeostasis, supporting and refining previously described interactions while posing novel signaling programs.

### Novel candidate regulators and localization by protein content of bronchiole-specific basal and secretory cell populations

At our finest level of subclass.L5 annotations, 69 clusters had molecularly distinct profiles, including 27 epithelial cell clusters (Fig. 6a). In Fig. 6b, we highlighted *TP63, KRT5, SFTPB, SCGB1A1*, and *SCGB3A2* expression as key molecular markers enabling characterization of conducting airway basal and secretory cells into *SFTPB^+^*bronchiole-specific subsets. Comparing Bronchial and Bronchiolar Epithelial spatial neighborhoods (Fig. 6c), *TP63^+^* basal cells were present in both. As expected, *SCGB1A1^+^* aw.secretory cells were more abundant in Bronchial Epithelial, and *SCBG3A2^+^* BrAS cells were largely seen in Bronchiolar Epithelial. *TP63, SCGB1A1*, and *SCGB3A2* positive populations each contained subtypes with and without *SFTPB* gene expression (Fig. 6d). In bronchi and bronchioles, SFTPB and

**Fig. 6:**
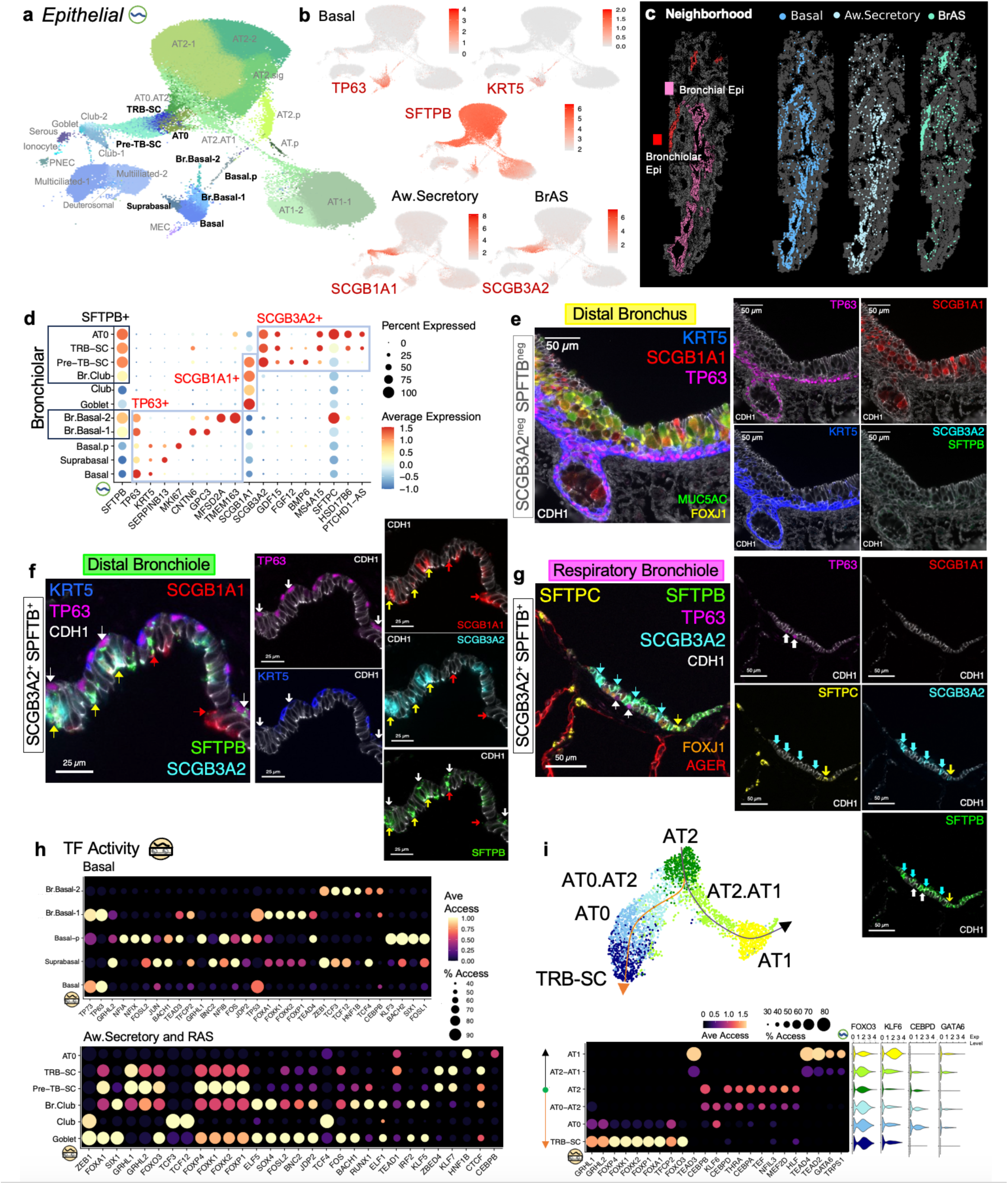
Multimodal characterization reveals bronchiolar-specific basal and secretory cells. **a**, UMAP embedding of subclass level 5 snRNA epithelial cells. **b**, Expression of key cell type and subtype defining markers for basal (TP63, KRT5), airway secretory (*SCGB1A1^+^* Aw.Secretory), and bronchiolar airway secretory (*SCGB3A2^+^* BrAS) cells. SFTPB expression is present in BrAS cells and a subset of basal and aw.secretory cells. **c**, MERFISH spatial maps colored by Bronchial and Bronchiolar Epithelial neighborhoods and individual maps of Basal, Aw.Secretory, and BrAS cells. Most BrAS cells are seen in the Bronchiolar neighborhood compared to Aw.Secretory cells which more present in the Bronchial neighborhood. Basal cells are seen in both Bronchial and Bronchiolar. **d**, Dot plot summarizing expression patterns of select airway epithelial cell subset-defining markers. *SFTPB* expression is seen in all bronchiolar airway epithelial cell subtypes. **e-g,** Multiplexed IF detection of airway-specific epithelial cells in (**e**) distal bronchus (SCGB3A2 and SFTPB negative), (**f**) distal bronchiole (SCGB3A2^+^SFTPB^+^), and (**g**) respiratory bronchiole (SCGB3A2^+^SFTPB^+^; SCGB1A1 and KRT5 negative). In each image, protein markers and corresponding color on display are listed accordingly. **f**, White arrows indicate TP63^+^SFTPB^+^KRT5^neg^ distal bronchiolar basal (Br.Basal) cells. Red arrows are SCGB1A1^+^ SFTPB^+^SCGB3A2^neg^ bronchiolar club (Br.Club) cells. Yellow arrows are SCGB3A2^+^SCGB1A1^+^SFTPB^+^ pre-terminal bronchiole secretory cells (pre-TB-SC). **g**, White arrows indicate distal Br.Basal. Turquoise arrows are SCGB3A2^+^SFTPB^+^SCGB1A1^neg^ TRB-SC. Yellow arrow indicates a SCGB3A2^+^SFTPB^+^SFTPC^+^ bronchiolar alveolar type 0 (AT0) cell. **h**, Dot plots of chromVAR average accessibility and proportion of cells accessible for top differentially active TFs for basal (top) and aw.secretory and BrAS (bottom) cell subtypes. Expression of TFs must be present in at least 10% of corresponding cell types. **i**, Slingshot trajectory, chromVAR top differentially active TFs, and expression of select TFs for TRB-SC, AT0, AT2, AT1, and intermediate alveolar subtypes. AT2 cells were designated as starting cell type.

SCGB3A2 protein localization began in primary bronchioles with progressively increasing expression noted distally in the terminal and respiratory bronchioles (Fig. 6e-g, Extended Data Fig. 9-10). Bronchi lacked both SFTPB and SCGB3A2 indicating that both serve as anatomic RNA and protein markers of varying bronchiolar-specific epithelial populations.

Our hierarchical annotation approach separated *SCGB3A2^+^*BrAS cells from *SCGB1A1^+^SCGB3A2^-^* aw.secretory cells (Fig. 6d). Aw.Secretory cells divided into Club and Goblet cell populations. A subset of *SCGB1A1^+^SCGB3A2^-^*club and *TP63^+^* basal cells were positive for *SFTPB^+^*which we localized and classified as bronchiolar club (Br.Club) and basal (Br.Basal) cells (Fig. 6e-g, Extended Data Fig. 9-10). In Fig. 6e, basal cells consistently positive for TP63 and KRT5, tightly lined the bronchial basal epithelial surface. In the bronchioles (Fig. 6f), *TP63^+^* basal cells were progressively more dispersed, and SFTPB co-staining was seen. In the respiratory bronchiole (Fig. 6g), KRT5 staining was negative, and only a few TP63^+^SFTPB^+^ basal cells were present. For *SCGB3A2^+^SFTPB^+^* BrAS cell subpopulations, *SCGB1A1^+^* pre-TB-SCs were identified in the distal bronchioles (Fig. 6f), *SCGB1A1^-^* TRB-SCs in terminal and respiratory bronchioles, and *SFTPC^+^* AT0 in respiratory bronchioles (Fig. 6g). Leveraging directly linked RNA and AC profiles to identify candidate TF regulators, Fig. 6h presents top differentially active TFs in subclass.L5 basal, aw.secretory and BrAS cells (Supplementary Tables 12 subclass.l5 DAR, enriched TFs, and active TFs). Candidate active TFs based on ChromVAR TF activity and expression in 10% of corresponding nuclei were notable for the following: TP63, FOXK1, FOXP1 for basal.SFTPB-1; HNF1B for basal.SFTPB-2; FOXA1, SIX1, SOX4, and KLF5 in goblet; ZEB1 and TCF4 for club-1; GRHL2, JDP2, RUNX1, and IRF2 for club-2; and GRHL1, FOXO3, FOXK1, and FOXP1 for pre-TB-SC.

Based on the prevailing model that AT2 cells serve as the resident alveolar progenitor capable of differentiating into AT1 cells during development and after injury, and informed by recent organoid models suggesting additional AT2 to TRB-SC differentiation^9,63,64^, we performed Slingshot trajectory analysis with AT2, AT2.AT1, AT1, TRB-SC, AT0, and AT0.AT2 cells (AT2 designated as the starting cluster). This pseudotime analysis recapitulated AT2 cell differentiation into AT1 cells and TRB-SCs with alignment of intermediate cell states along expected trajectory paths (Fig. 6i). Differential active TF analysis revealed that during AT2 cell differentiation, intermediate states retained expression of AT2-specific TFs but exhibited decreased accessibility of corresponding TF binding sites. For example, in the AT2 to AT1 cell trajectory, AT2.AT1 cells expressed similar levels of *CEBPD* but lacked enrichment for CEBPD TF binding sites compared to AT2 cells. AT2.AT1 cells began to demonstrate activity (accessibility and expression) for AT1-specific TFs such as GATA6. During the transition from AT2 to TRB-SCs, AT0.AT2 and AT0 cells retained *CEBPD* expression however, AT0 cells showed reduced CEBPD accessibility and gained TF activity similar to TRB-SCs. AT0.AT2 and AT0 cells also gained KLF6 activity. Our dual-omic trajectory suggested that AT2 cells proceed through intermediate states by closing accessible regions for AT2-specific TFs and incrementally opening TFBS for cell type reprogramming. Overall, matched SNARE2-AC and RNA profiles enabled the identification of novel TF candidates that regulate basal, aw.secretory, and BrAS cell subtypes. Spatial transcriptomics and proteomics validated molecular and spatial localization of these key populations. Trajectory analysis further predicts sequential TF shifts that drive cell type differentiation.

## Discussion

Lung scRNA-seq studies have redefined concepts of lung cell plasticity, regionalization of injury and repair, and therapeutic discovery. Examples include *TP63^+^KRT17^+^KRT5^-^*basal-like or aberrant basaloid cells and *SCGB3A2^+^* distal airway specific secretory cells which have been identified in IPF, COPD, and other advanced parenchymal lung diseases, with a suggestion of interrupted cell type transitions in an effort to repair^6,8,9,22,65,66^. *SCGB3A2^+^SFTPB^+^*distal airway cells as respiratory airway secretory (RAS) cells^6^, pre-terminal bronchiole secretory cells (pre-TB-SC), terminal and respiratory bronchiole secretory cells (TRB-SC), and TRB-specific alveolar type-0 (AT0) cells along with terminal airway-enriched secretory cells (TASC)^8,9^. As SCGB3A2 and SFTPB gene/protein expression is seen proximal to terminal and respiratory airways but is exclusive to bronchioles, we propose bronchiolar airway secretory cells (BrAS) as a unifying, anatomy-informed collective term for SCGB3A2^+^SFTPB^+^ populations. The emergence of these disease-associated cell types underscores the need to investigate their origins, mechanisms of transformation, and therapeutic relevance. Integrating spatial and regulatory data into cell- and network-level analyses will promote disease modeling. Due to fibrosis and loss of cellular integrity, diseased lungs are often more amenable to single-nucleus-based multiomic profiling. As such, a comprehensive lung single-nuclear reference is important for aligning physiologic cell types and detecting pathologic cell states. To address this need, we present a multimodal nucleus-centric resource that confirms and expands ‘omic characterization of known and novel molecular profiles, including bronchiolar-specific epithelial cell subsets along with proliferative, progenitor, and intermediate states relevant to homeostasis, disease, and regeneration. As single cell/nucleus datasets expand, harmonizing ‘omic profiles across modalities and adopting a unified, anatomy-informed nomenclature will be essential to enable cross-study comparisons and accelerate translational discovery.

In support of a unified framework, we present a tiered dictionary (Supplementary Tables 4-5) of healthy adult human lung cells and marker genes, applying consensus cell names aligned with the latest cell ontology terms. We deliver companion multilevel FAIR NS-Forest cell type markers to serve as minimum criteria to harmonize nomenclature and guide spatial ‘omic panel design. Complementing transcriptomic profiles, we add a gene regulatory layer capturing precise AC regions using SNARE-seq2. Candidate active TFs, expressed in setting of corresponding open binding sites, align with previous studies and highlight novel TF regulators, including for bronchiole-specific epithelial cells, intermediate/transitional, and proliferative states. To our knowledge, this snHLA resource contains the largest paired human adult lung AC and transcriptomic library to date.

Focusing on clinically relevant bronchioles, the multimodal snHLA enables high-resolution classification of epithelial cell types with aligned transcriptomic, accessible chromatin, and spatial RNA and protein signatures. Our anatomically targeted approach complements the resolution achieved with technically difficult microdissection of human distal airways, providing support for consensus nomenclature for SCGB3A2^+^SFTPB^+^ cells. We substantiate the presence of bronchiolar basal and secretory populations with MxIF spatial localization, add new SCGB1A1^+^SFTPB^+^SCGB3A2^-^ Br.Club and SCGB3A2^+^SFTPB^+^SFTPC^+^ABCA3^+^ AT0.AT2 profiles, and contribute regulatory elements and candidate TF drivers for each. In silico, *SCGB3A2^+^* cells have been implicated as the cell of origin for *TP63^+^KRT17^+^KRT5^-^* aberrant basaloid cells in IPF^22^. Likewise, distal airway basal cells were the region-specific cellular origin for TASC after acute lung injury in vitro^8^. Collectively, bronchiole-specific basal and secretory cell plasticity are compelling targets to resolve IPF and related diseases. Newly identified regulator targets will facilitate deciphering mechanisms causing alterations in key lung silent zones, regions of lung disease initiation.

Beyond reference tables, histologic images, and expert pathologist review, all data and images are accessible in an interactive format on the HuBMAP portal (portal.hubmapconsortium.org). This open access, spatially resolved, multimodal transcriptomic, regulatory, and protein atlas, developed in collaboration with the NIH BRINDL tissue repository (brindl.urmc.rochester.edu), offers an unprecedented resource. By enabling direct access to both richly annotated datasets and the corresponding primary biospecimens, this platform empowers the research community to explore, apply, and expand upon the snHLA. Together, these tools open new avenues to address critical gaps in lung development, health, aging, and disease.

## Methods

### Human Tissue Processing and Nuclei Isolation

All human lung samples were obtained from BRINDL (https://brindl.urmc.rochester.edu) supported by the NIH Common Fund HuBMAP and the NHLBI LungMAP programs and overseen by the University of Rochester (IRB approval # RSRB00047606). Donor metadata is listed in Supplementary Table 1. Organ acceptance from the Organ Procurement and Transplantation Network (OPTN), tissue acquisition, processing and storage protocols can be found on <u>protocols.io</u>^67–69^. For 10X snRNA-seq and SNARE-seq2, approximately 1 cm^3^ blocks of mapped, OCT embedded, frozen lung tissue were utilized. From each block, six to ten 40 µm serial section curls were combined for nuclear isolation. A 5-10 µm section before and after approximately every 10th section was collected to document histology. One frozen section from each block used was stained with hematoxylin and eosin (Supplementary Fig. 1) and reviewed by pathologist (G.D.). Due to the nature of frozen unfixed lung tissue, some sample drop-out was unavoidable in the thin H&E. Five-micron sections of formalin fixed paraffin embed tissue blocks were prepared and used for MxIF^70^. Representative FFPE sections were reviewed by pathologist for structures and pathology. One H&E per case are demonstrated in Supplementary Fig. 2.

### Single Nucleus Omic Assays and Data Processing

10X snRNA-seq and SNARE-seq2 dual RNA and ATAC-sequencing^14^, quality control, clustering, and downstream analysis was performed as previously described^71^. Briefly, 10X snRNA sequencing was accomplished using 10X Chromium Single-Cell 3’ Reagent Kits v3. Applying the 10X Cell Ranger v3 pipeline with the GRCh38 (hg38) reference genome, sample demultiplexing, barcode processing, and gene expression quantifications including introns were completed. For SNARE-seq2, AC and RNA libraries were sequenced on the Illumina NovaSeq 6000 system using the 300 cycle and 200 cycle reagent kits, respectively. An automated pipeline was applied (github.com/huqiwen0313/snarePip) for quality assessment, doublet removal, cell clustering, peak generation, and differential accessible region identification.

For 10X snRNA-seq quality control, cell barcodes passing 10X Cell Ranger filters were used for downstream analyses. Mitochondrial transcripts (MT-*) and doublets identified using the DoubletDetection software (github.com/JonathanShor/DoubletDetection, v2.4.0)^72^ were removed. All samples were combined across experiments and cell barcodes having greater than 200 and less than 7500 genes detected were kept.

For SNARE2-RNA, cell barcodes for each sample were retained with the following criteria: DropEst (github.com/kharchenkolab/dropEst)^73^ cell score greater than 0.9; greater than 200 UMI detected; greater than 200 and less than 7500 genes detected. To further remove low quality datasets for snRNA-seq and SNARE-seq2 RNA, a gene UMI ratio filter (gene.vs.molecule.cell.filter) was applied using pagoda2 (github.com/kharchenkolab/pagoda2)^74^. Then for SNARE2-RNA, doublets identified by both DoubletDetection (v3.0)^72^ and Scrublet (github.com/swolock/scrublet, v0.2.2)^75^ were removed. For SNARE2-AC, cell barcodes which already passed SNARE2-RNA quality filtering were further retained with the following criteria: transcription start site (TSS) enrichment greater than 0.15; at least 1000 read fragments and at least 500 UMI; and fragments overlapping the promoter region ratio of greater than 0.15.

### Clustering and Cell Type Annotation

Clustering of snRNA was performed using pagoda2. Counts were normalized to the total number per nucleus, batch variations were corrected by scaling expression of each gene to the dataset-wide average. After variance normalization, all 4937 significantly variant genes were used for principal component analysis. Clustering was performed at different k values (50, 100, 200, 500) based on the top 50 principal components and the infomap community detection algorithm. Then, principal components and infomap cluster annotations for each k value were imported into Seurat (v4.0.0)^76–79^ for subsequent analyses. Uniform manifold approximation and projection (UMAP) dimensional reduction was performed using the top 50 pagoda2 principal components. For each infomap cluster k resolution, differentially expressed genes between all clusters were identified using the Seurat FindAllMarkers function (only.pos = TRUE, max.cells.per.ident = 5000, logfc.threshold = 0.25, min.pct = 0.25). The primary cluster resolution (k = 500) was chosen based on the extent of clustering observed. Then, a cluster decision tree was implemented to determine whether a cluster should be split further. Clusters with distinct markers at lower k values were kept. If clusters at lower k values occurred across major classes (epithelial, endothelial, mesenchymal, immune), the clusters were labeled as low quality and removed. Two subsequent rounds of iterative clustering were completed to identify and remove low quality cells.

Using consensus annotations from LungMAP CellCards as a foundational guide^15^, cluster annotation was based on positive and negative cell type markers. Proliferating cell subsets were identified by expression of cell cycle S and G2/M phase genes (e.g. MKI67, TOP2A, and CIT). To further resolve airway epithelial cells, non-alveolar epithelial clusters and neuronal like cells were re-clustered as described above (2392 significantly variant genes). Clusters with distinct markers at k=50 were kept. Where possible and to the best of our ability, supplemental references were used to annotate clusters with consistent gene signatures (see Supplementary Table 4). Overall, 98 clusters (k500mod) from the cluster decision tree were retained for annotation.

Cluster annotations were also organized under class and subclass level categories based on increasing granularity using a hierarchical approach which incorporated lung cell compartment or lineage, anatomical structures, and increasing cluster resolution (lower k values). For example, in subclass level 4 and 5, lower k clusters with distinct gene signatures were annotated as a numbered subset within a higher-level subclass annotation, e.g. AT2-2 (see Supplementary Table 4).

### Integrating 10X snRNA-seq and SNARE-seq2

The snRNA data was down sampled to 2,000 cells max per cluster prior to integration. Similar to previous^71^, integration of 10X snRNA-seq and SNARE2 RNA data was completed using Seurat (v4.0.0) with snRNA as the reference. Counts were normalized (SCTransform), anchors were identified based on snRNA pagoda2 principal components (FindTransferAnchors), and SNARE2 data was projected onto the snRNA UMAP embedding (MapQuery). Integrated clustering was done using pagoda2 and integrated principal components. Integrated clusters were annotated by the most overlapping, correlated and/or predicted snRNA cluster annotation. Annotations were confirmed by manual inspection of cell type markers. The addition of Multicil.Aw.Sec intermediate cells were identified. Clusters that overlapped with different classes of cell types were labeled as low quality and removed. To visualize the entire snRNA and SNARE2 datasets together, a joint UMAP embedding was generated using Sketch to subsample 50,000 representative cells from each assay. Then, transfer anchors were identified using snRNA sketch as reference. ProjectData extended SNARE2 sketch cells and full datasets onto the snRNA sketch UMAP embedding.

### NS-Forest Cell Type-Specific Marker Genes

To identify a minimal set of characterizing marker genes that can identify cell types annotated at each subclass level from the clustering analysis of the 10X snRNA-seq data, the validated NS-Forest method^20^ – a random forest machine learning-based method that determines the cell type-specific necessary and sufficient marker gene combinations from sc/snRNA-seq experiments – was used on a dataset downsampled to 5000 cells max per subclass.l5 annotations, ensuring a balanced representation of cell type diversity at subclass level l5. NS-Forest v4.0 algorithm (<u>github.com/JCVenterInstitute/NSForest</u>) was applied with the “gene_selection = ‘BinaryFirst_high’” setting to pre-select candidate genes with high binary expression patterns, i.e., highly expressed in the target cell type and little to no expression in the off-target clusters, as input to the random forest step of the algorithm. The NS-Forest algorithm produces four performance metrics – F-beta score, precision, recall, and On-Target Fraction – to quantify the cell type classification accuracy and marker gene expression specificity for each cell type. The NS-Forest results, including the NS-Forest marker genes, top 10 binary genes, and performance metrics for all subclass levels are in Supplementary Table 4 and 6. At subclass.L1, 22 unique markers were identified to characterize the 12 subclass.L1 cell types with median F-beta score of 0.75 and median precision of 0.95, indicating very strong classification performance. At subclass.L5, 142 unique markers were identified to characterize the 69 subclass.L5 cell types with median F-beta score of 0.60 and median precision of 0.77, also suggesting strong classification performance given the number of diverse cell types at this level.

### RNA Correlation Analysis

Using available reference datasets^2,4^, correlation analysis was performed as described previously. From the integrated Human Lung Cell Atlas (HLCA) core^4^, nasal and tracheal cells were excluded based on their common coordinate framework anatomical location (CCF score = 0, 0.36). Briefly, a set of common variable genes were determined through the intersection of variable features identified for each reference and our snRNA dataset. Average expression was calculated using these intersected variable genes. Average expressions between reference datasets and snRNA were correlated and plotted using the corrplot package^80^.

### SNARE2-AC Analysis

All SNARE2 nuclei barcodes passing both RNA and subsequent ATAC QC filtering were retained. We applied the following SNARE-AC QC thresholds: >500 fragments per nuclei, transcription start site (TSS) enrichment > 0.15 & fraction reads in promoter (FRiP) > 0.15). Peak calling was performed separately for k500mod clusters, subclass.L3, subclass.L4, and subclass.L5 annotations using MACS (github.com/macs3-project/MACS, v3.0.0)^81^ CallPeaks function. We then used Signac (github.com/stuart-lab/signac, v1.1.1)^82^ reduce followed by FeatureMatrix and CreateChromatinAssay to generate a peak count matrix and Seurat object, respectively. Jaspar motifs (JASPAR2024 Core, Homo Sapians)^83^ were used to generate a motif matrix and motif object.

### Differentially Accessible Regions (DARs) and TF analysis

DARs for subclass.L3-5 were identified using CalcDiffAccess function (github.com/yanwu2014/chromfunks)^84^ which accounts for technical differences in cell total accessibility. P-values were calculated using a Fisher’s Exact Test on a hypergeometric distribution and adjusted p-values (or q-values) were calculated using the Benjamini & Hochberg method. Subclass cell annotations with >10 nuclei/cluster, >100 DARs, p-value < 0.01, and log fold change > 1 were visualized using SWNE ggHeat^84^. DAR accessibility or peak counts were averaged and scaled (trimming values to a minimum of 0 and a maximum of 4). Motif enrichment for cell type DARs was completed using Signac FindMotifs hypergeometric test. To identify top expressed cell-type-specific TFs, ChromVAR TFBS activity scores were calculated using Signac RunChromVAR^82,87^. Differential activities of TFBS were assessed using the Seurat FindAllMarkers (min.pct = 0.05, logfc.threshold = 0.1, max.cells.per.ident = 5000, mean.fxn = rowMeans). Then, only differentially active (p value < 0.05) TFs with expression in at least 10% of nuclei (snRNA) within the corresponding subclass clusters were included. Top ChromVAR TFBS of expressed TFs (i.e., active TFs) between clusters were visualized using Seurat DotPlot.

### Cell-cell Communication Analysis

In the snRNA dataset, clusters with <200 cells were removed and remaining subclass.L3 clusters were down sampled to 1,000 cells each. Cellchat v2^58^ was applied using default parameters and the Cellchat human ligand-receptor database. When identifying over-expressed ligands, receptors, and ligand-receptor interactions using computeCommunProb function, we set population size = FALSE to reduce the overpowering signaling strength of abundant cell populations. Additional Cellchat analysis outputs for different subclass levels and when applying a subset of the human database with only secreted signals are available on our Github.

### Gene Expression, AC and TF Figures

All UMAP, feature, dot, and violin plots for 10X snRNA-seq and SNARE2 data were generated using Seurat. Correlation plots were generated using the corrplot package. *Genome coverage plots were performed using Signac.

### Pseudo-time Trajectory Analysis

Slingshot pseudo-time trajectory analysis was performed on subset of distal epithelial cells as previously described^22^, including evaluation of trajectory robustness. Loess plots for DEGs of interest were made using ggplot2 geom_smooth on the Slingshot trajectory variable timeline^88^.

### Spatial Cell-type Verification

### Multiplexed Immunofluorescence (MxIF) Chemistry Assay and Data collection

Identities and locations for cell types at subclass level 3 to 4 by protein content were sought by an approach based on the CODEX MxIF staining technology^89^. The protocol used is published at protocols.io^90^. In brief, blocks (up to 1.5 x3 cm) of mapped, inflation-fixed, FFPE lung tissue were sectioned (5 micron) and mounted on TruBond 380 glass slides, or in one case on a slide coated with indium-tin oxide (ITO) for subsequent MALDI-Mass spec analysis. After drying at 60^0^ overnight, the slides were deparaffinized in xylene, cleared in 100% ethanol, and rehydrated via a decreasing ethanol series prior to High pH <u>H</u>eat-Induced <u>E</u>pitope <u>R</u>etrieval (HIER) (see protocol). After cooling, lung tissue was washed with hydration buffer (Akoya Biosciences, Marlborough, MA) and then incubated for 20-30 min in staining buffer (Akoya). Sections were then covered with 200 µL of antibody buffer composed of staining buffer supplemented with recommended concentrations of N, G, J, & S blockers (Akoya) and the specified dilutions of up to 45 antibody-barcode conjugates (Supplementary Table 7) followed by incubation in a humidified chamber at room temperature for 3 h. Sections were then washed with staining buffer followed by fixation in 1.6% paraformaldehyde for 10 min. After a series of rinses in PBS, samples were placed in a storage buffer (Akoya) and photobleached by illumination with a 200 mA, 15 watts, 1600 lumens bulb overnight at 4°C. Image acquisition was performed using the Phenocycler-Fusion™ 2.0 platform utilizing the 20X (0.5 micron/pixel) objective^90^. Image processing was automated via Fusion 2.2.0 software according to manufacturer recommendations. Acquired .qptiff files were converted to ome.tiff for upload to local OMERO imaging database (www.openmicroscopy.org) utilizing bfconvert (openmicroscopy.org/bio-formats/downloads/). Figures were generated in Omero.figure. Phenoimager images reported are subsequently displayed in the HuBMAP portal. Further details on MxIF panels can be found in the HuBMAP Organ Mapping Panels at (<u>humanatlas.io/omap</u>). Validation of antibody staining is reported in Antibody Verification Reports (avr.hubmapconsortium.org/). Validation was performed by Pathologist (G.D.) and subject matter experts, and is based on commercial antibody reports of specificity, as well as comparison to prior publications, prior experience and reported specific protein staining in the Human Protein Atlas (proteinatlas.org).

### Multiplexed Error-robust Fluorescence In Situ Hybridization (MERFISH)

Frozen, non-fixed, OCT embedded tissue block samples were prepared according to Vizgen’s MERFISH protocol (Vizgen Doc. number 91600002) with some deviations as reported in protocols.io^91^. Samples were cryosectioned to 10μm thickness, kept at -20°C for 2 hours to encourage adherence to the treated coverglass, prior to fixation at 47°C with pre-warmed 4% (vol/vol) paraformaldehyde (PFA) in 1xPBS for 30 min. Next, 50μL of MERSCOPE 500 Gene Panel Mix: Panel ID VA86 was added directly on top of the sample. Sections were hybridized for 48 hours in a humidified chamber at 37°C. Samples were then gel embedded and treated with Vizgen’s Digestion Mix for 2 hours at room temperature (RT). Further cell clearing was achieved with an incubation in Clearing Solution for 16 hours at 47°C in a humidified chamber, followed by an incubation of approximately 24 hours at 37°C interrupted by 3 hours of autofluorescence quenching in a MERSCOPE Photobleacher at RT. A MERSCOPE 500 Gene Imaging Cartridge was activated, and the sample was loaded into the MERSCOPE Flow Chamber. Regions were selected from the mosaic with a focus on capturing as many cells from the tissue as possible.

### MERFISH Data Analysis

For gene panel selection and probe design, we selected 503 genes for MERFISH imaging on the Vizgen MERSCOPE platform. Our selection of target genes was based on multiple strategies, which included, but were not limited to: 1) cell type marker genes; 2) genes as ligands, receptors, or transcription factors that may play a role in human lung maturation and cell differentiation; 3) genes encoding ECM molecules; and 4) genes detected in a distributed subdomain of a cellular cluster on a UMAP, suggestive of heterogeneity of the cluster. To confirm that the genes selected are suitable for cell type identification, we re-clustered published snRNA-seq data using only the selected genes to confirm that the cell types could be identified^92^. The final gene panel consisted of 498 genes and 5 additional sequential (smFISH) genes after some were determined by Vizgen to be un-targetable by MERFISH probes. The code library and probe design were optimized by Vizgen; see Supplementary Table 10 for final library and probes.

Transcripts were called as part of the MERSCOPE’s built in analysis pipeline using the MERlin package^93^. Segmentation was performed on the DAPI stain using Cellpose (v.2)^94^ ‘cyto-2’ model^95^. This model provides about one-micron of nuclear expansion for nuclear segmentation. Once new cell boundaries were identified, transcripts were assigned to these cells and a new cell by gene table was generated. Next, the cell-by-gene table was filtered to remove cells with total counts < 10, as well as cells with a volume below 4 times the median volume. The data was then normalized (target sum per cell; 10,000).

snRNA-seq data was used as reference data to perform cell type annotation in MERFISH by integrating the two datasets and performing label transfer, similar to methods seen in previous publications^96,97^. Prior to integration, snRNA subclass.L3 cells types without markers present in the MERFISH gene panel (MEC, neutrophil, neural) and those not expected to have distinct markers (RAS.AT2, AT2.AT1) were removed. The snRNA-seq data and MERFISH datasets were each subset to only include the 496 overlapping genes found in both datasets. Both datasets were then normalized and combined into one object, followed by PCA (components = 50) and integration using Harmony^98^. A k-nearest-neighbors classifier was trained using the Harmony adjusted snRNA-seq reference data and asked to predict the classification of the cells in the MERFISH data. This classifier then assigned the subclass.L3 labels to the MERFISH cells based on their nearest neighbor in the Harmony PCA embedding, and these labels were used for further MERFISH analysis (hence, referred to as “Harmony annotations”). To identify mesothelial cells, we extracted a distinct cluster of cells satisfying the following marker gene criteria: WT1 >= 1 count, GAS1 >= 1 count, UPK3B >= 3 count. Due to difficulty in differentiating between select cell types with the necessarily limited MERFISH probeset, closely related cell types were combined (e.g. ASMC and VSMC into SMC). Rare ionocytes and PNECs were assigned to a higher proportion of cells than expected and produced non-characteristic spatial patterns, and thus were removed prior to the final Harmony integration and label transfer with snRNA-seq and MERFISH.

### MERFISH Neighborhood Analysis

To generate spatially resolved clusters from MERFISH data, we employed the BANKSY algorithm^57^. A spatial neighbor graph was constructed using 50 nearest neighbors (k_geom = 50), with a scaled Gaussian function applied as the neighbor weight decay to emphasize local spatial relationships. The algorithm incorporated both the mean expression, and the azimuthal Fourier transform matrix of gene expression within each cell’s local neighborhood, capturing both average expression levels and spatial patterns. We set the mixing parameter λ to 0.999, which places significant emphasis on neighborhood features in the clustering process. A λ value close to 1 shifts the focus toward domain segmentation by heavily weighting spatial context, whereas a value of 0 would be equivalent to non-spatial clustering based solely on global gene expression profiles. Dimensionality reduction was performed using PCA, retaining 20 principal components for clustering. Subsequently, the Leiden algorithm was applied to assign cells to clusters based on the reduced-dimensional space, using a resolution parameter of 0.6.

## Dataset Availability

https://portal.hubmapconsortium.org/browse/publication/a10041ad9ebae0b42d3c7f602ba37b82

## Code Availability

All analysis was completed using published R packages.

## Acknowledgements

We are extremely grateful for the generosity of the Donor families; we honor their loss and their selfless contribution to research. This work was supported by the Departments of Pediatrics at the University of California San Diego, La Jolla, CA, University of Rochester, Rochester, NY, the University of North Carolina, Chapel Hill, NC, the Department of Bioengineering, University of California San Diego, La Jolla, CA, the Department of Cellular and Molecular Medicine, University of California San Diego, La Jolla, CA, the Department of Pathology, Seattle Children’s Research Center, Seattle, WA, and the Division of Intramural Research (DIR) of the National Library of Medicine (NLM), National Institutes of Health, Bethesda, MD with funding awards from the National Heart, Lung, and Blood Institute (NHLBI) Molecular Atlas of Lung Development Program Human Tissue Core (LungMAP HTC) grants U01HL122700 and U01HL148861 (to GSP), the NIH Common Fund grant U54 HL165443-01 (to GSP, with Co-Investigators TED, GD, JH, XS, and KZ) and 1R03OD036499-01 (to YZ), U54 HL145608 (to KZ, XS, JH), 5K12HD105271-04 (to TED), the Chan Zuckerberg Institute (CZI) Human Lung Cell Atlas 1.0 (to TED), and NIH 1S10OD032391-01 (to QZ). The data used for bulk RNA-seq correlation with MERFISH described in this manuscript were obtained from: the GTEx Portal dbGaP Accession phs000424.v10.p2 on 07/23/2025. The Genotype-Tissue Expression (GTEx) Project was supported by the Common Fund of the Office of the Director of the National Institutes of Health, and by NCI, NHGRI, NHLBI, NIDA, NIMH, and NINDS. Bioinformatics infrastructure in this publication was supported by the University of Rochester Clinical and Translational Science Award (CTSA) (UL1 TR002001). We gratefully recognize many supporting personnel including bioinformatics support from Jeanne Holden-Wiltse (University of Rochester Medical Center, URMC), Anthony Corbett (URMC), Jennifer Dutra (URMC). We acknowledge the support and contributions of the National Institution of Health Program Staff and our co-investigators in the Human Biomolecular Atlas Program Consortium.

## Author Contributions

TED, JH, XS, KZ and GSP conceived the project and the overall design of the experimental strategy. GSP, RM, HH, JV, and GD contributed to tissue collection, pathology review, and processing. TED, DD and KC conducted 10X snRNA-seq and SNARE-seq2 experiments. JMP generated and curated MxIF data. ZZ performed MERFISH experiments. TED and IB analyzed 10X snRNA-seq and SNARE-seq2 data. BP and YZ performed NS-Forest analysis. EB, JO, and SP analyzed MERFISH data. JMP led generation of MxIF figures with assistance from TED and GSP. JV, RM, ZB, YZ, RS, QZ, GD, JH, XS, KZ, and GSP provided critical intellectual input and data interpretation. TED and GSP drafted the majority of the manuscript with input and review from all authors.

## Author Information

Dinh Diep, Kimberly Conklin, and Kun Zhang

Present address: San Diego Institute of Science, Altos Labs, San Diego, CA, USA

