## Extended Data Figures 1-10 for "A Multimodal Spatial and Epigenomic Atlas of Human Adult Lung Topography"

**Extended Data Fig. 1: Anatomic characterization of healthy adult human lung donor tissue and multimodal RNA comparison.**

**a**

| ASSAY | No. of blocks | DONOR (metadata and no. of blocks) | | | | | | | | |
|---|---|---|---|---|---|---|---|---|---|---|
| | | Sex | | Age Group | | | Race | | | |
| | | F | M | Young 19-25y | Middle 30s | Older 49-59y | White | Black | Asian | Hispanic |
| 10x snRNA-seq | 44 | 4 | 7 | 5 | 2 | 4 | 6 | 2 | 1 | 2 |
| SNARE-seq2 | 27 | 4 | 3 | 3 | 2 | 2 | 2 | 2 | 1 | 2 |

**b**

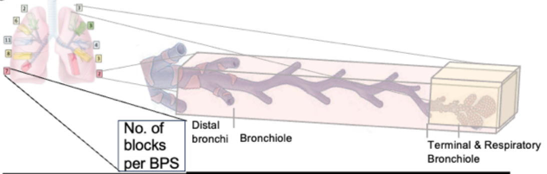


| Right Lung | | | No. of blocks by largest airway captured | | | | |
|---|---|---|---|---|---|---|---|
| Lobe | Bronchopulmonary Segment (BPS) | | Distal Bronchi | Bronchioles | | | |
| | | | | Primary | Lobular | Terminal | Respiratory |
| Superior | Apical | 2 | 1 | | 1 | | |
| | Posterior | 6 | 2 | | 2 | 2 | |
| | Anterior | 11 | 3 | | 7 | | 1 |
| Middle | Lateral | 8 | 3 | | 3 | 2 | |
| | Medial | 7 | 2 | 1 | 2 | 2 | |
| | TOTAL | 34 | 11 | 1 | 15 | 6 | |

| Left Lung | | | No. of blocks by largest airway captured | | | | |
|---|---|---|---|---|---|---|---|
| Lobe | Bronchopulmonary Segment | | Distal Bronchi | Bronchioles | | | |
| | | | | Primary | Lobular | Terminal | Respiratory |
| Superior | Apicoposterior I | 3 | 1 | | 1 | 1 | |
| | Apicoposterior II | 3 | 1 | 1 | 1 | | |
| | Anterior | 4 | 2 | | | 2 | |
| | Superior lingular | 3 | 3 | | | | |
| | Inferior lingular | 2 | 1 | | 1 | | |
| | TOTAL | 15 | 8 | 1 | 3 | 3 | |

**c**

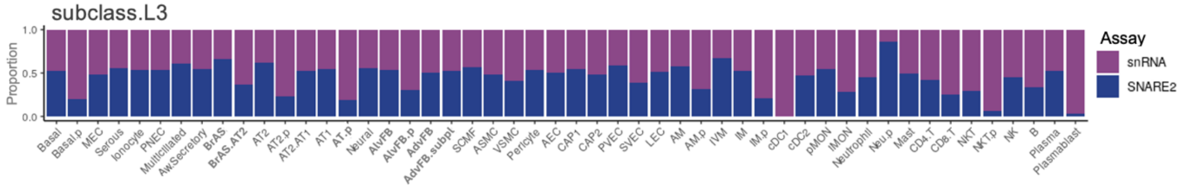


**d**

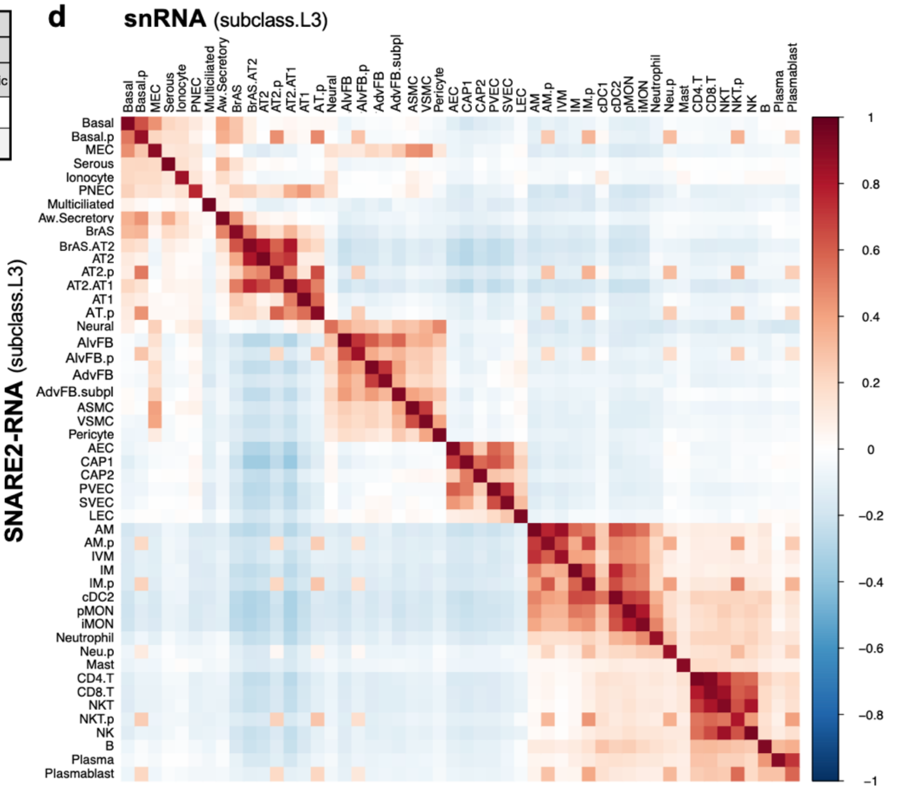


**a**, Metadata table for lung blocks processed for each single-nucleus assay. 11 total donors were profiled. N = 22 lung blocks with both snRNA and SNARE-seq2 data, 22 blocks with snRNA-seq only, and 5 blocks with SNARE-seq2 only. **b**, Tables summarize number of deceased human donor lung blocks sampled by lung lobe, bronchopulmonary segment (BPS), and largest distal airway structure captured. Based on histology and cell types captured, we assumed that each block sampled contains all subsequent distal airway structures. Diagram above the tables provide examples of distal bronchi and terminal bronchiole lung blocks. See Supplementary Tables 1 and 2 for details. **c**, Proportion of cell type nuclei captured per assay for harmonized subclass.L3 annotations. **d,** Heatmap with correlation of averaged scaled gene expression values for snRNA-seq and SNARE-seq2 for subclass.L3. Off center correlations are largely due to similarities in proliferating cell types called.

**Extended Data Fig 2: Quality control assessment of 10X snRNA-seq atlas.**

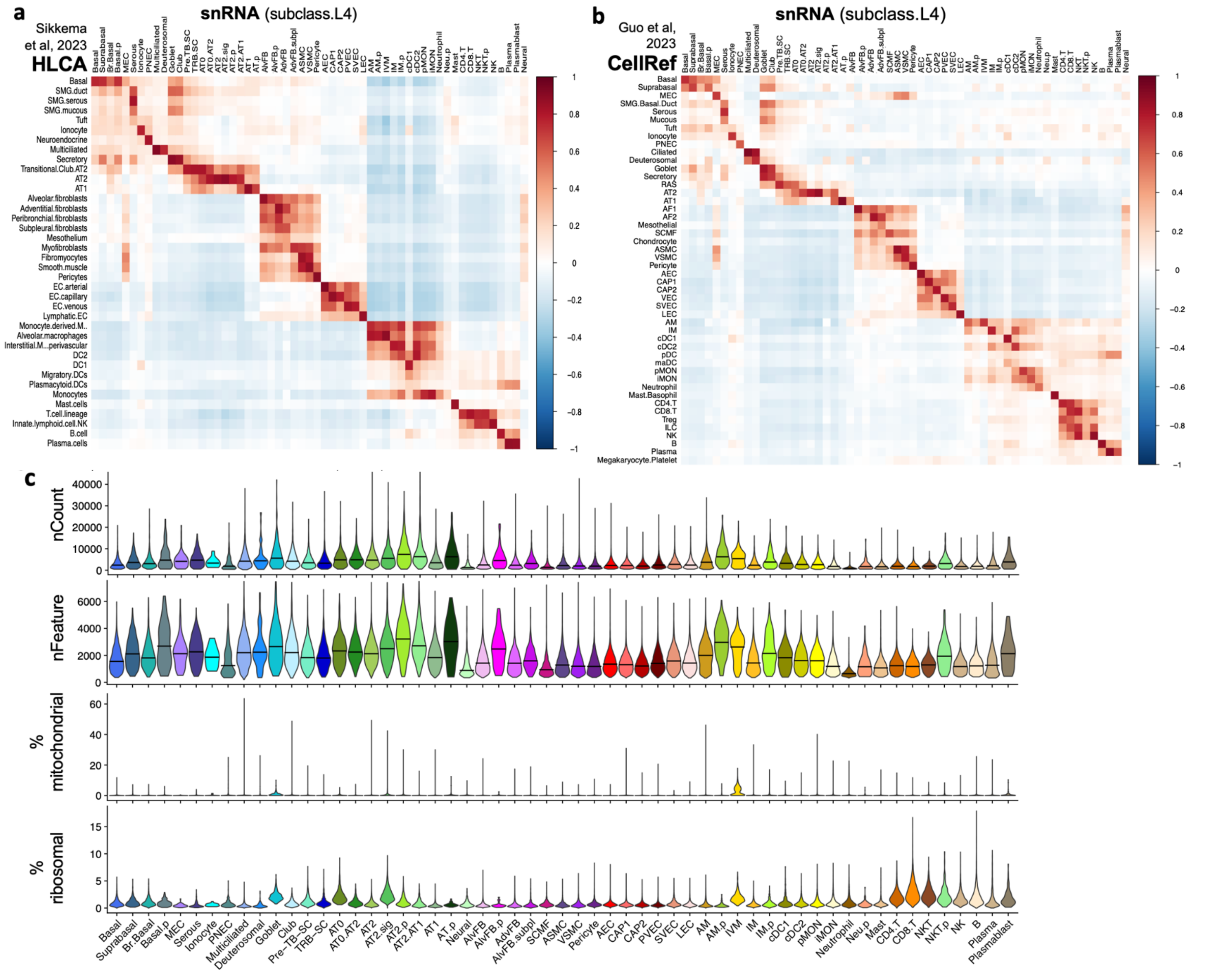


**a-b,** RNA correlation of snRNA subclass.L4 annotations and published lung atlases (Sikkema et al., 2023 and Guo et al., 2023). **c,** Violin plots for read counts, genes (features), percent mitochondrial reads, and percent ribosomal reads for snRNA subclass.L4 cell types. See also Supplementary Table 3.

**Extended Data Fig. 3: Spatial proteomics localizes MKI67+ proliferating adult lung cell populations.**

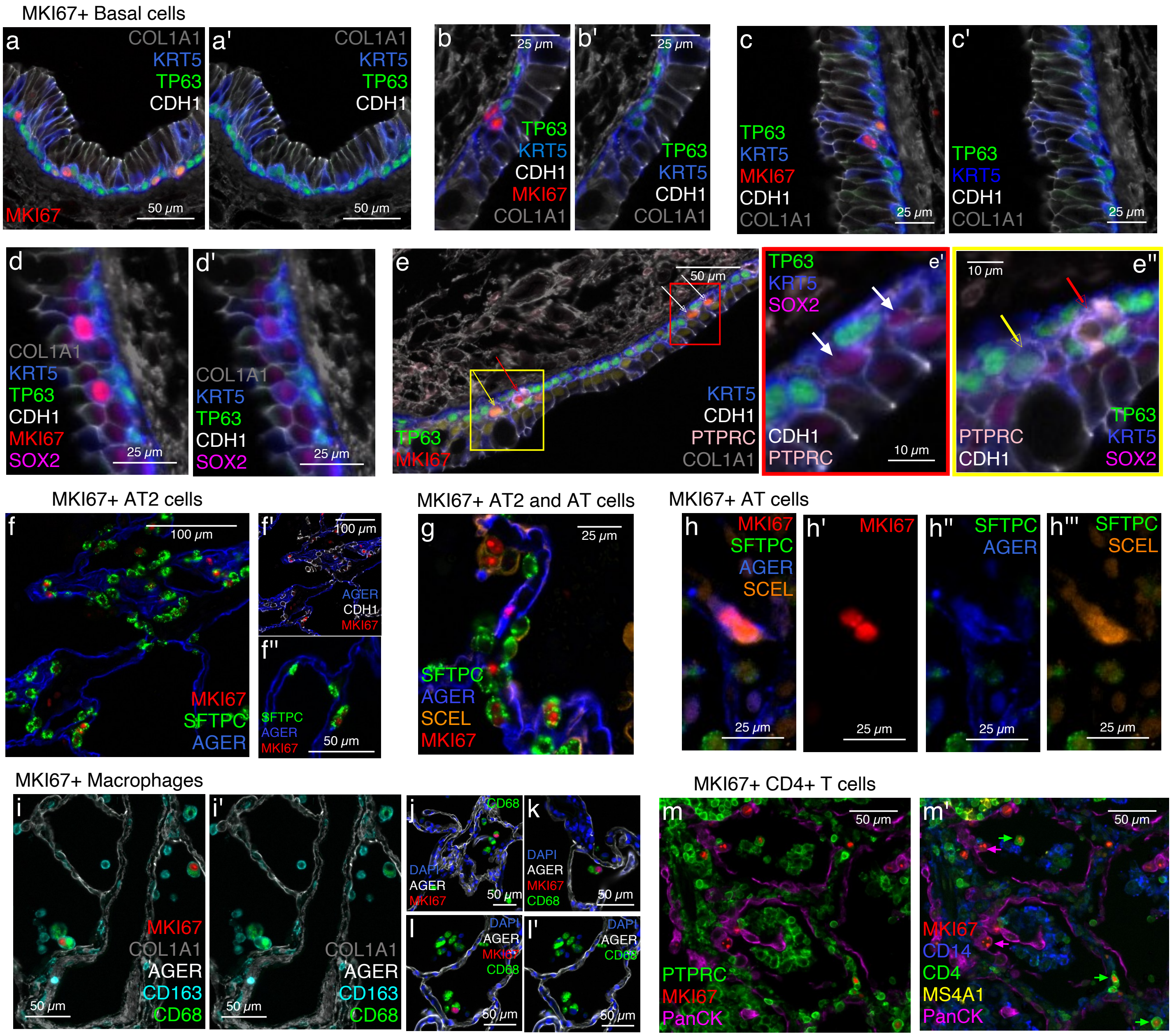


**a-d**, MKI67+ nuclei in TP63+KRT5+ proliferating basal (basal.p) cells in three bronchial airways. **a'-c'**, Same fields of view without MKI67 to demonstrate that while the MKI67+ basal cells are KRT5+, TP63 protein appears reduced in proliferating or differentiating cells (Li et al., Oncogene 2023). **d-d'**, Demonstrates MKI67+ cells that are SOX2+KRT5+, minimal TP63+, in near basal position of airway epithelium. **e-e''**, Demonstrates variation in identity of MKI67+ cells in basal layer: **e'** red box, indeterminate airway epithelial cell, SOX2+, white arrows; **e''** yellow box, TP63+KRT5+, yellow arrow and PTPRC/CD45+ leukocyte, red arrow. **f**, Several MKI67+SFTPC+ proliferating AT2 (AT2.p) cells in the alveolar septum. **f'**, Same field of view with SFTPC removed confirming MKI67+ epithelial cells (CDH1+, white). **f''**, High power view of MKI67+ nucleus within surfactant protein of AT2 cell. **g**, Demonstrates MKI67+ nuclei in the alveolar septal wall surrounded by cytoplasm that is either SFTPC+ or AGER+ or SCEL+ consistent with cycling alveolar epithelium of various cytotypes. **h-h'''**, MKI67+ alveolar transitioning epithelial cell (AT.p, SFTPC negative, AGER+, SCEL+). **i-l**, CD68+ macrophages are detected mostly in alveolar space. **i'** and **l'** with MKI67 off to demonstrate CD68+ cytoplasm. **m-m'**, MKI67+ cells tend to occur more frequently in areas of leukocyte aggregation in the alveolar region, some are PTPRC+CD4+ (green in respective panels) suggesting activated tissue resident memory T cells, others are cytokeratin (PanCK, purple) positive epithelial cells. Four lung cases are represented in the figure (D260, D273, D292, D346).

**Extended Data Fig. 4: Spatial proteomics resolves adult lung leukocyte aggregates.**

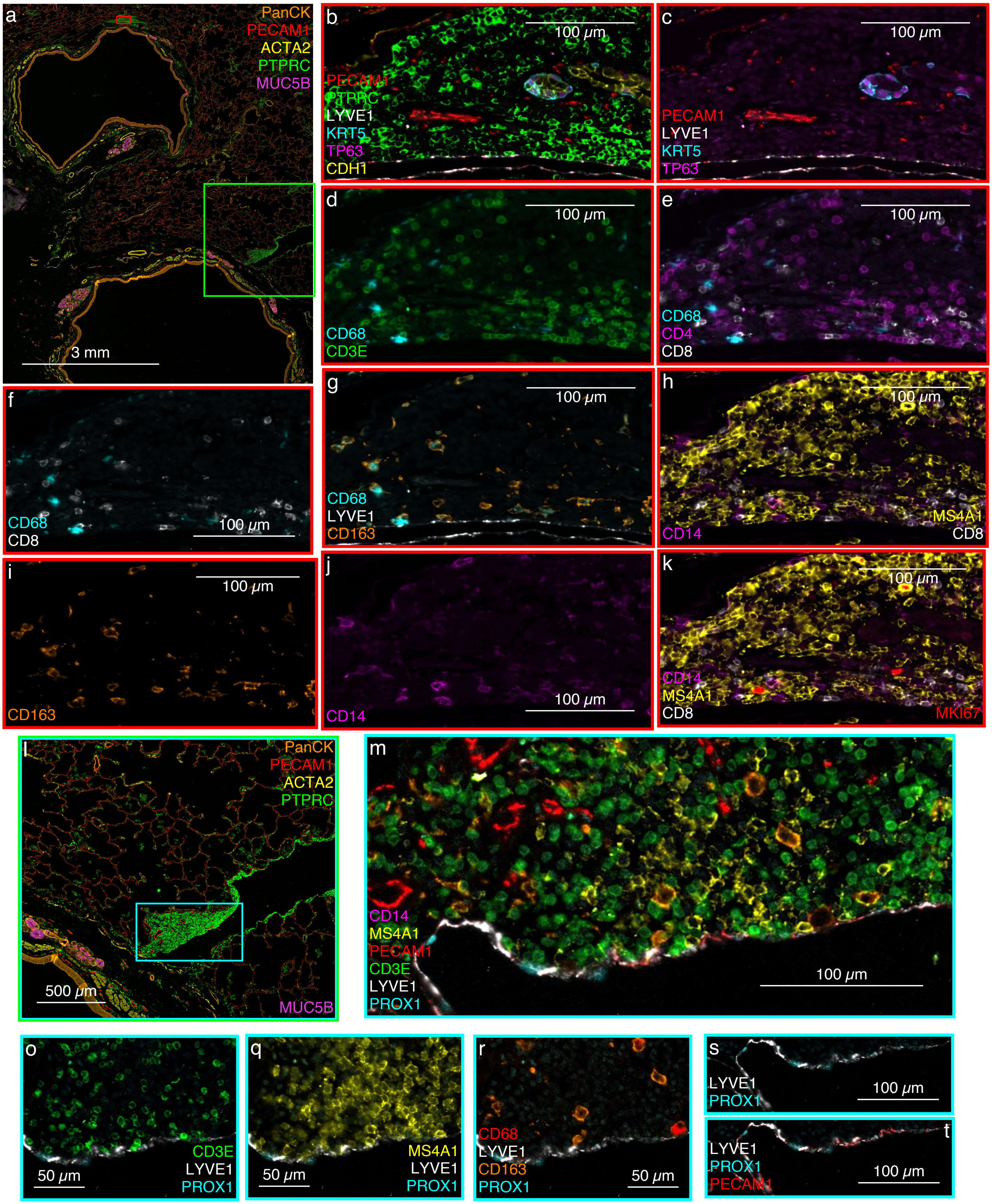

Representative multiplexed immunofluorescent (MxIF) images in adult lung. **a**, Two distal bronchi (I.D. 2-5 mm) with airway epithelium (pan-cytokeratin, PanCK, orange), smooth muscle bands (ACTA2, yellow), vasculature (PECAM1, red), submucosal glands (MUC5B, purple) and leukocyte aggregates (PTPRC, CD45, green) in bronchial submucosa (red box) and peri-lymphatic vessel (green box). **b-k**, Red box inset from **a**. **b-c**, Highlight cell type markers in vascularized, bronchus associated lymphoid tissue, including leukocytes (PTPRC, green) and blood vessels (PECAM1, red), bordering a lymphatic vessel (LYVE1, white), with central epithelial cell structures (CDH1, yellow) consistent with submucosal gland duct basal cells (KRT5, turquoise, TP63 nuclei, pink) and ductal epithelium (CDH1, yellow). **d-k**, Highlight immune cells including CD4+ (pink) or CD8+ (white) T lymphocytes (CD3, green), CD68+ (turquoise) +/- CD163 (orange) tissue macrophages, a B cell predominant zone (MS4A1, CD20, yellow), and tissue monocytes (CD14, purple in **h, k**). **k**, Demonstrates MKI67+ nuclei, three consistent with MS4A1+ B cells and one undefined. **l-t**, Details of vascularized leukocyte aggregate (PTPRC, green) near bronchus from green box inset in **a**. **m-t,** This aggregate of leukocytes neighbors a lymphatic vessel (LYVE1, white, PROX1, turquoise) and is composed of CD3E+ T lymphocytes (green), CD20+ B lymphocytes (MS4A1, yellow), and CD68+ (red) or CD163+ (orange) tissue macrophages/monocytes.

**Extended Data Fig. 5: MERFISH quality control assessment.**

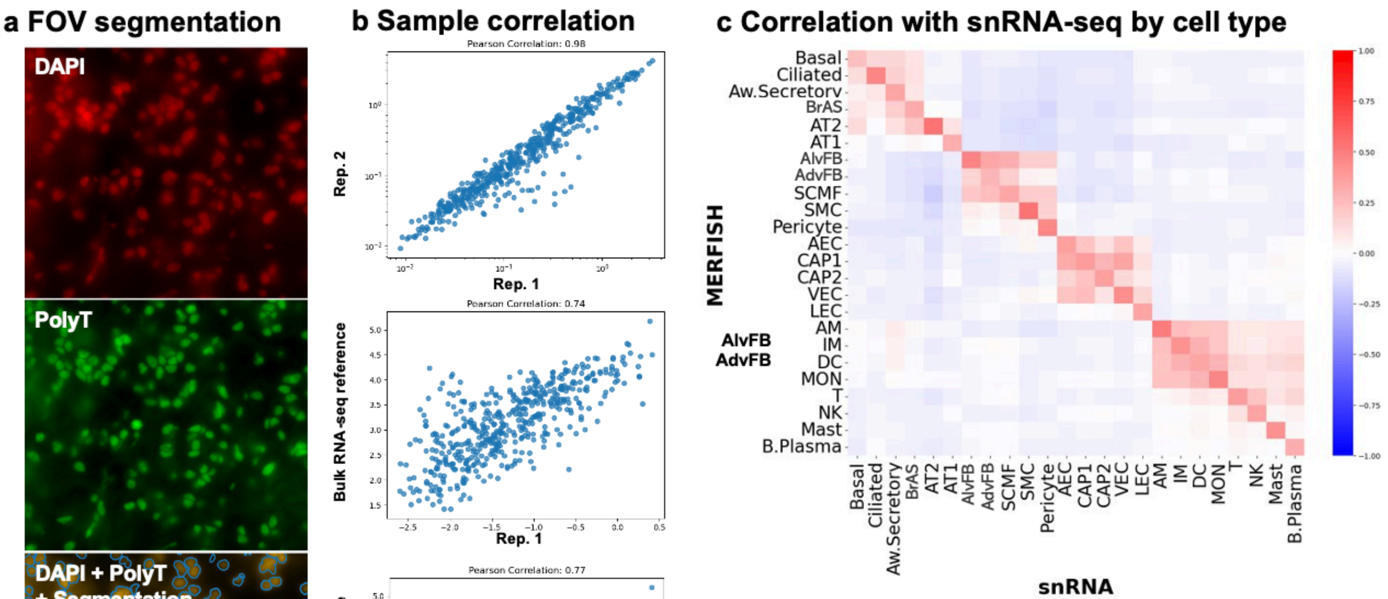


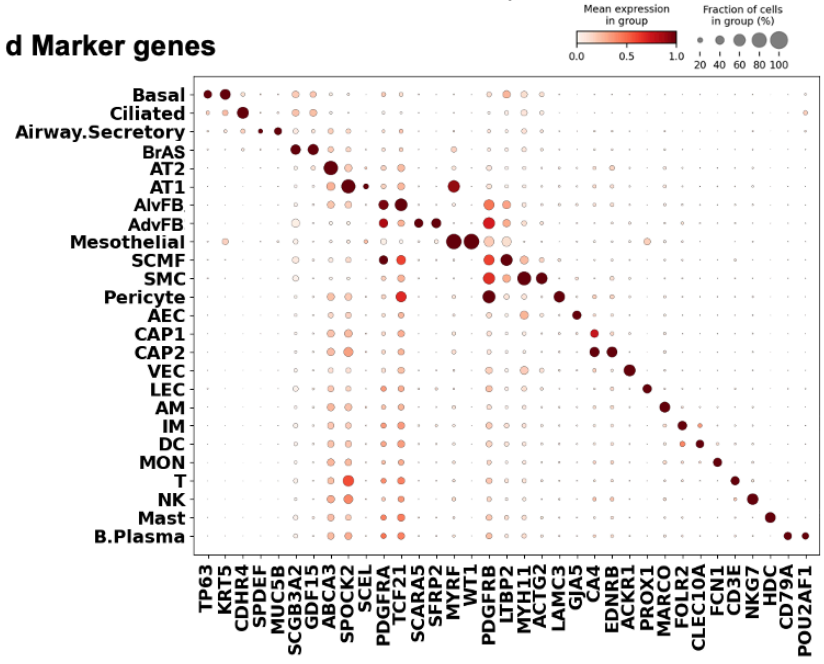


**e Cell type counts by assay**

| | # of cells scRNA | # of cells MERFISH | % of cells scRNA | % of cells MERFISH |
|---|---|---|---|---|
| Basal | 5042 | 2307 | 1.5 | 0.89 |
| Ciliated | 10444 | 7624 | 3.11 | 2.93 |
| Aw.Secretory | 2213 | 2151 | 0.66 | 0.83 |
| BrAS | 4670 | 3010 | 1.39 | 1.16 |
| AT2 | 50879 | 43656 | 15.17 | 16.77 |
| AT1 | 28035 | 24230 | 8.36 | 9.31 |
| AlvFB | 31505 | 30172 | 9.4 | 11.59 |
| AdvFB | 7304 | 9723 | 2.18 | 3.74 |
| SCMF | 1489 | 1736 | 0.44 | 0.67 |
| SMC | 9668 | 13803 | 2.88 | 5.3 |
| Pericyte | 5854 | 15120 | 1.75 | 5.81 |
| AEC | 9953 | 5189 | 2.97 | 1.99 |
| CAP1 | 51709 | 24453 | 15.42 | 9.4 |
| CAP2 | 19442 | 17054 | 5.8 | 6.55 |
| VEC | 13272 | 7270 | 3.96 | 2.79 |
| LEC | 4220 | 4983 | 1.26 | 1.91 |
| AM | 21709 | 13810 | 6.47 | 5.31 |
| IM | 7897 | 3927 | 2.36 | 1.51 |
| DC | 3569 | 2581 | 1.06 | 0.99 |
| MON | 11770 | 6242 | 3.51 | 2.4 |
| T | 20100 | 10269 | 5.99 | 3.95 |
| NK | 7636 | 5268 | 2.28 | 2.02 |
| Mast | 3469 | 1969 | 1.03 | 0.76 |
| B.Plasma | 3438 | 3599 | 1.03 | 1.38 |
| Mesothelial | 0 | 126 | 0 | 0.05 |

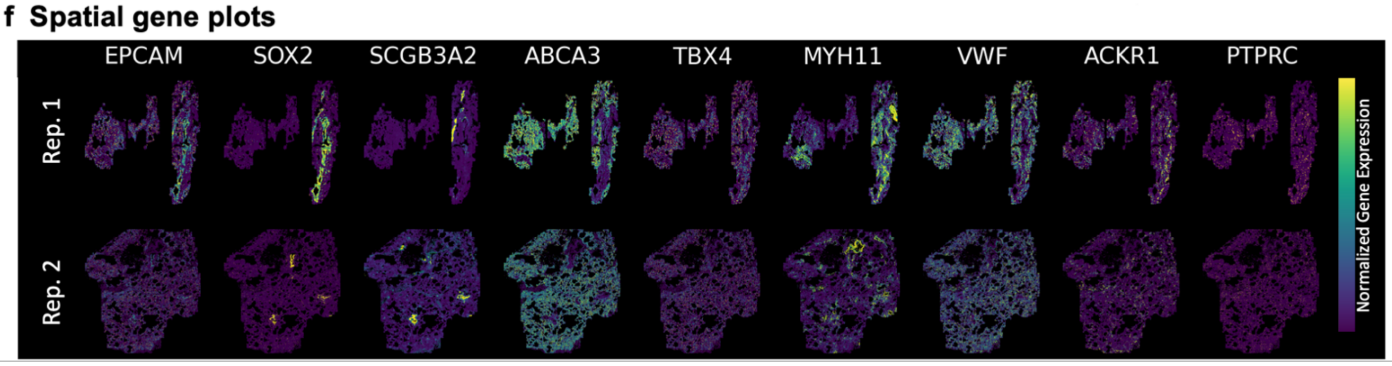

**a**, Single field of view (FOV) from MERSCOPE with two stains and final segmentation. **b**, Correlation of MERFISH experiments with each other and Genotype-Tissue Expression (GTEx) Project bulk lung reference RNA-seq data. **c**, Correlation of MERFISH data with snRNA by assigned cell types. **d**, Prevalence of a selection of marker genes in each assigned cell type. **e**, Total cell or nuclei counts and percentage of each cell type from MERFISH experiments and snRNA-seq. **f**, Spatial gene plots for select genes with distinct and expected regions of expression.

**Extended Data Fig. 6: Spatial communities identified by BANKSY and zone analysis.**

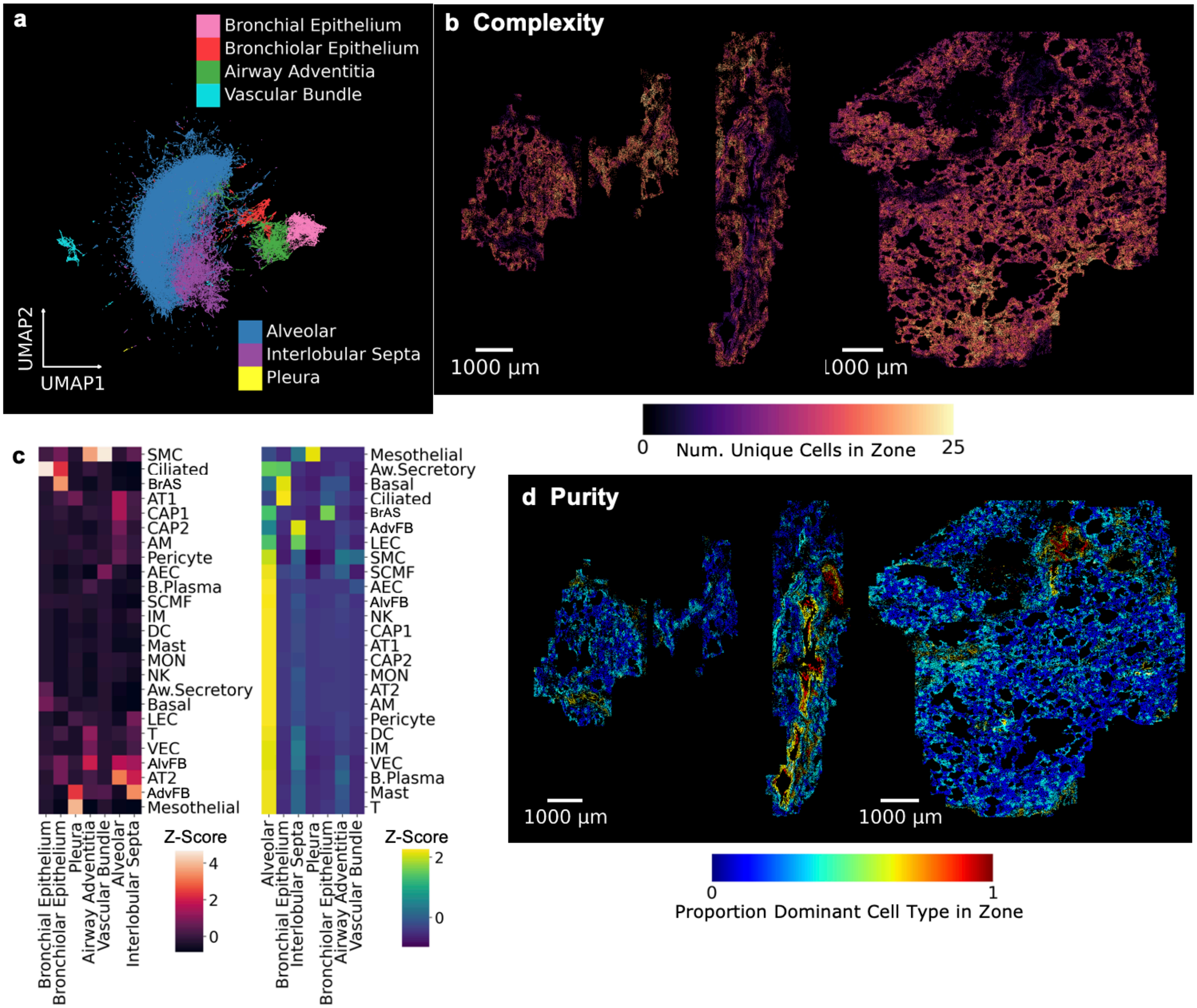


**a**, BANKSY clustering UMAP projection of cells colored by collapsed and named BANKSY clusters (lung structural neighborhoods). **b**, Spatial map of cells colored by Complexity. Complexity is defined as the number of unique cell types in a zone around the cell. **c**, Clustered heatmaps showing the relation between cell types and collapsed BANKSY clusters. The left heatmap is Z-scored across BANKSY clusters, reflecting relative cell type counts to the spatial community cluster. The right heatmap is Z-scored across cell types, reflecting relative spatial community counts to the cell type. **d**, Spatial map of cells colored by Purity. Purity is defined as the proportion of the dominant cell type in a zone around the cell.

**Extended Data Fig. 7: Global and major cell-cell signaling pathways in human adult lung.**

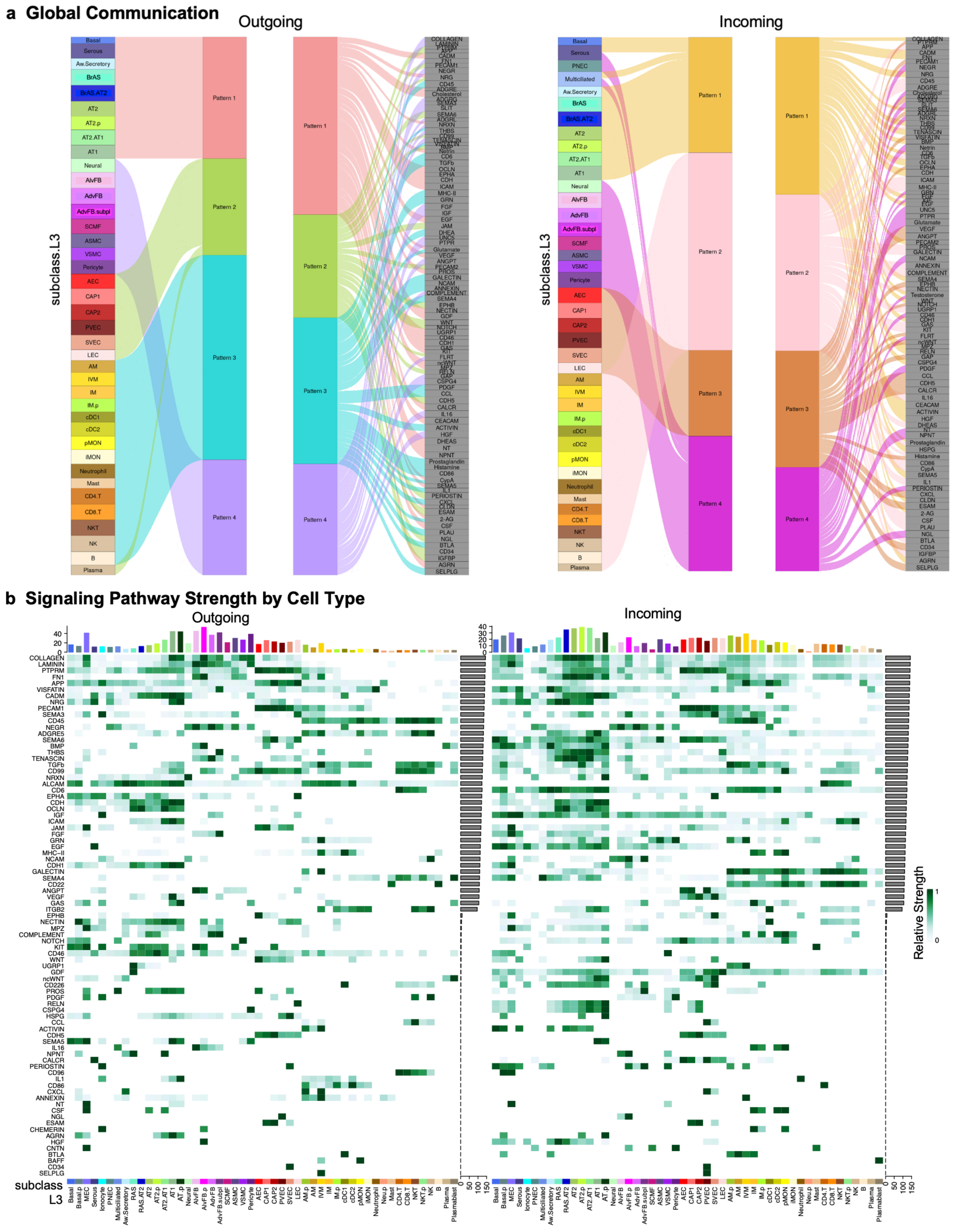

**a**, Pattern recognition of global coordinated signaling identified 4 outgoing (left) and 4 incoming (right) patterns. Heatmaps illustrate contribution of lung cell types (subclass.L3) to each pattern and enriched signaling pathways in each pattern. See also Supplementary Table 18. **b**, Heatmaps of relative strength of outgoing (left) and incoming (right) signaling pathways for all lung cell types at subclass.L3. Y-axis gray bars represent row-scaled relative signaling strength for each pathway across all cell types. Top X-axis colored bars demonstrate total signaling strength for each cell type across all signaling pathways.

**Extended Data Fig. 8: Divergent signaling pathways in alveolar and adventitial fibroblasts.**

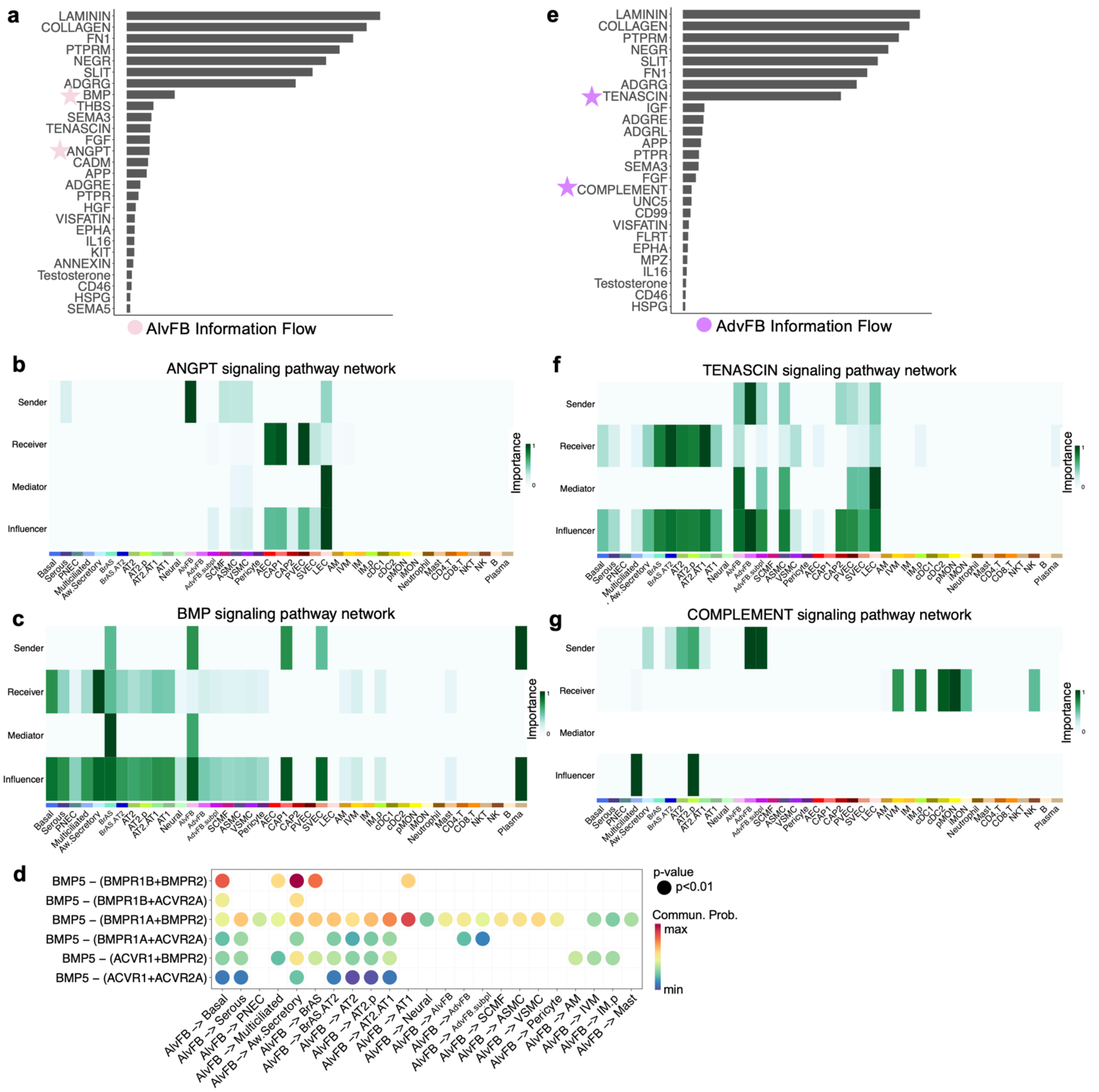


Bar plots rank top pathways for AlvFB (**a**) and AdvFB (**e**) cells serving as the signaling source. Stars indicate signaling pathways highlighted in panel **b-c** and **f-g**. **d**, Dot plot of BMP pathway communication probability between AlvFB (sender) and cell type specific receivers.

**Extended Data Fig. 9: Spatial proteomics delineates airway specific markers for bronchiolar epithelial subsets.**

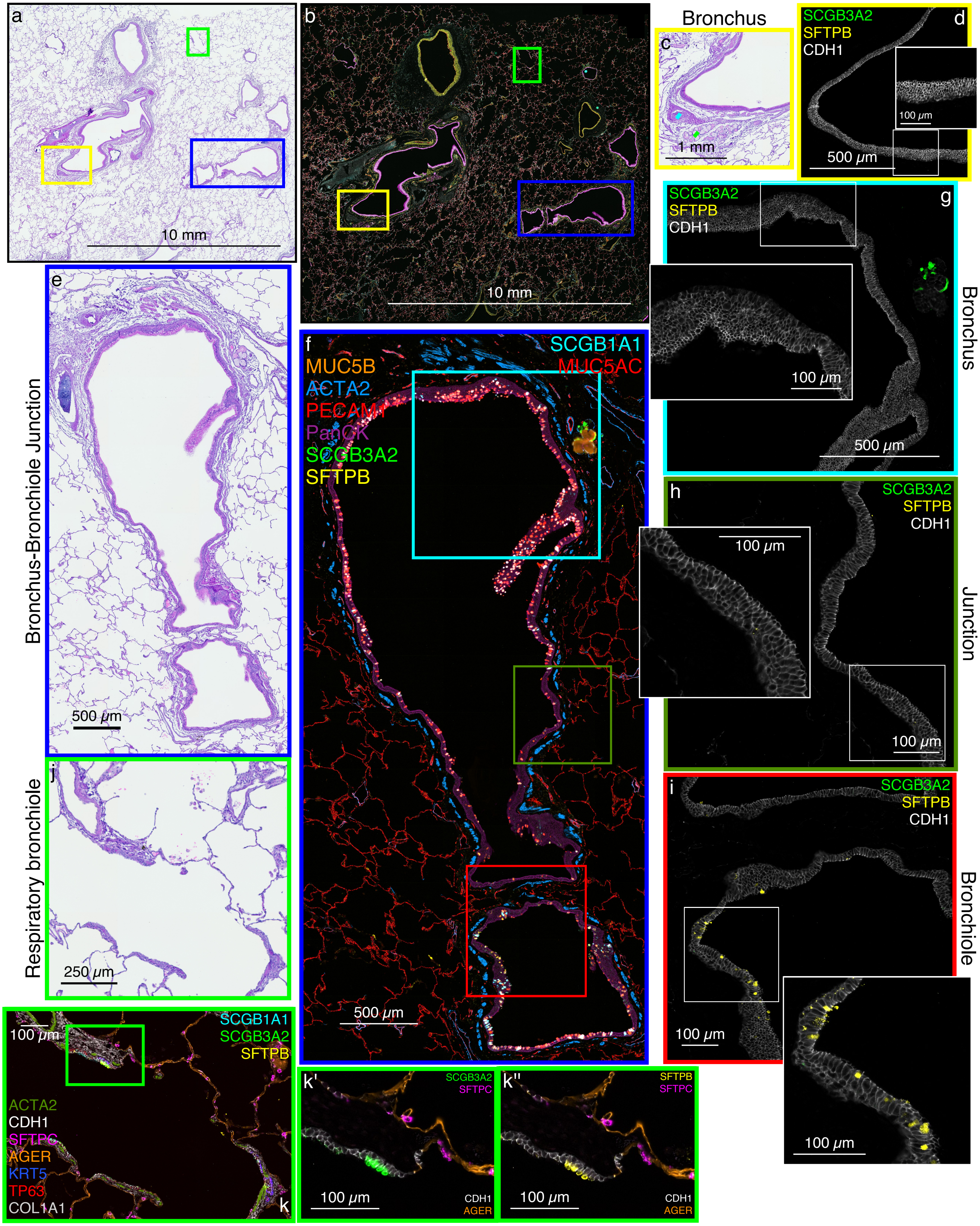

Multiplexed immunofluorescence (MxIF) delineates relative location of key bronchiolar-specific markers by protein content. CDH1 (white) discretely outlines airway epithelium and is used throughout for cell segmentation. **a-b**, H&E stain and MxIF of same tissue section (donor D341) with boxed regions of interest (ROI). **c-d**, Distal bronchus (panel **a** yellow ROI) identified by submucosal glands (MUC5B+ SMG, green diamond in H&E) and cartilage plate (blue diamond in H&E). SCGB3A2 (green) is present only in SMG (not shown) and SFTPB is not detected in bronchi, confirmed by presentation of select markers in **d** with accompanying inset. **e-f**. H&E and MxIF of longitudinal section of airway bordered by ACTA2+ smooth muscle (blue in MxIF) from upper bronchus (cyan ROI) through transition (green ROI) to proximal bronchiole (red ROI). **g**, Bronchial epithelium with undetectable SFTPB and SCGB3A2; **h**, Epithelium at junction of bronchus to bronchiole with rare SFTPB detected; **i**, Proximal bronchiolar epithelium begins to show SFTPB+ cells. **j-k**, H&E and MxIF of respiratory bronchiolar epithelium with prominent SCGB3A2+SFTPB+ cells.

**Extended Data Fig. 10: Spatial proteomics maps progression of airway-specific basal and secretory cells from small bronchi to respiratory bronchioles.**

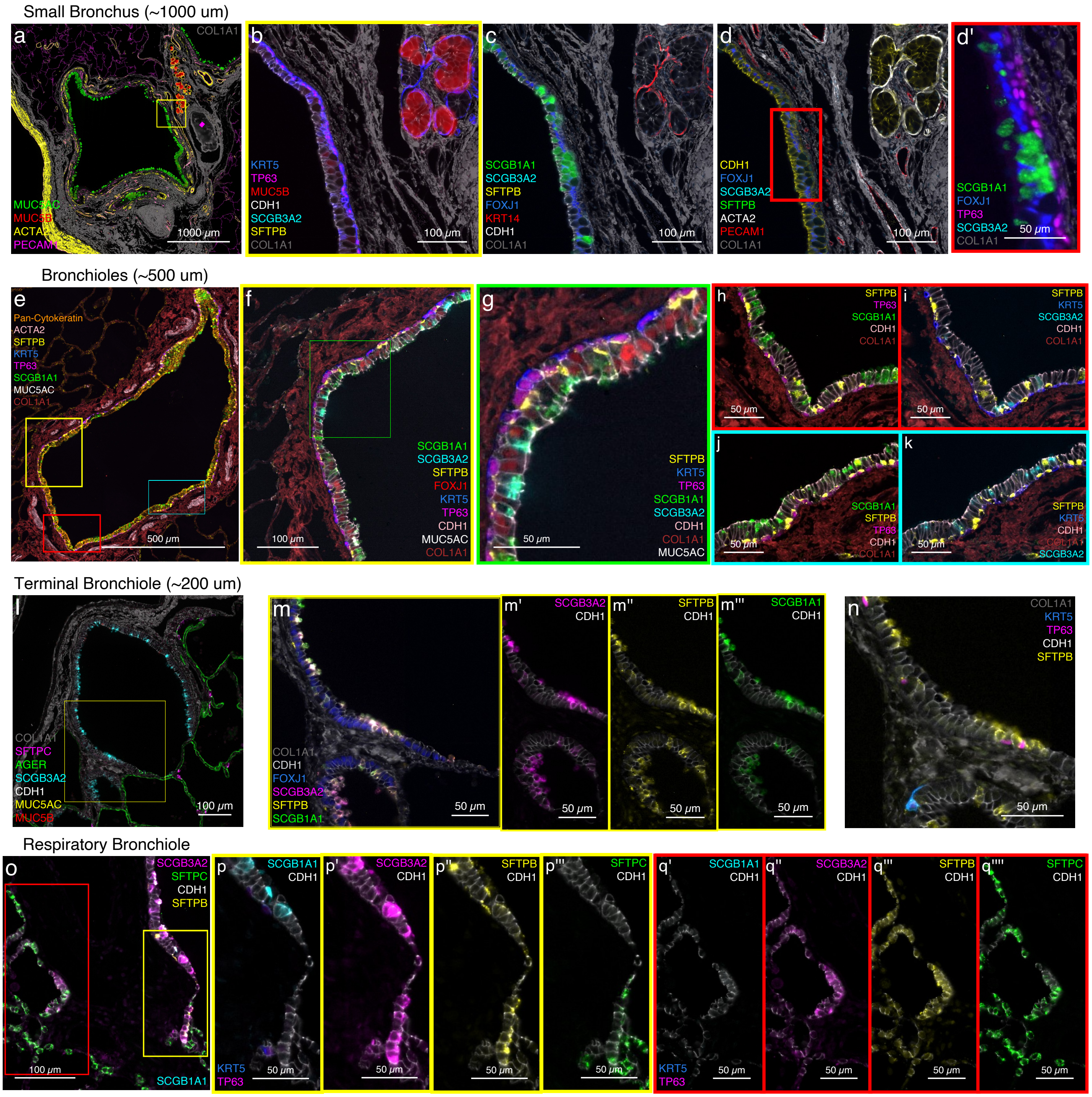

**a**, Bronchus (I.D. ~1000 μm) identified by submucosal glands (SMG, MUC5B, red), bundles of smooth muscle (ACTA2, yellow), submucosal vasculature (PECAM1, pink and ACTA2), cartilaginous plates (pink diamond), and position within extracellular matrix (ECM including COL1A1, grey) next to ACTA2+ continuous smooth muscle of a pulmonary artery (lower left). Rich mucin production includes MUC5AC and MUC5B. **b**, A layer of basal cells with cytoplasmic KRT5 and nuclear TP63 line the airway. Both are also detected in SMG myoepithelial cells along with KRT14 and ACTA2 in panels **c** and **d**. MUC5B staining is strong in SMG mucus cells and present but weaker in the goblet cells of the airway epithelium. **c**, SCGB1A1 is present in both MUC5B positive and negative cells. **d** and **d'**, FOXJ1+ multiciliated cells are interspersed with SCGB1A1+ aw.secretory cells. No SFTPB nor SCGB3A2 is detected in the bronchial epithelium. **e-k**, Medium sized bronchioles (I.D. 500-1000 μm) are characterized by little or no mucin proteins (minimal MUC5AC, white, **e-g**), decreased KRT5, persistent SCGB1A1, and appearance of SCGB3A2 and SFTPB. SCGB1A1 is detected independent of SFTPB and SCGB3A2 (Club) but also co-localizes with SFTPB alone (Br.Club), and with both SCGB3A2 and SFTPB (Pre-TB-SC). **g, h-i, and j-k**, In addition to SCGB1A1+ aw.secretory subtypes, three representative regions of the small bronchiole epithelium demonstrate TP63+ basal cells with both KRT5 (blue) and SFTPB (yellow) and mutually exclusive KRT5 (blue with purple nucleus, basal) or SFTPB (yellow with purple nucleus, distal Br.Basal). SFTPB is occasionally noted at the apical surface of secretory cells with little or no SCGB1A1 or SCGB3A2. FOXJ1 (nuclear, red) multiciliated cells form the majority of the small bronchiolar epithelium. Occasional cells of the epithelium are not well defined by this antibody panel. **l-m**, Terminal bronchioles (I.D. ~200 μm) now contain terminal and respiratory bronchiole secretory cells (TRB-SC) co-expressing SCGB3A2 and SFTPB (negative for SCGB1A1, MUC5AC, and MUC5B) in the apical cytoplasm. **n**, TP63+ basal cells are infrequent in the terminal bronchioles, are often SFTPB+ and rarely KRT5+. **o-q**, Respiratory bronchiole epithelium is continuous with alveolar epithelium and includes TRB-SC cells (SCGB3A2+, SFTPB+, SFTPC-, SCGB1A1-) and AT0 cells (SCGB3A2+, SFTPB+, SFTPC+, SCGB1A1-).
