## Supplementary Figures 1-4 for "A Multimodal Spatial and Epigenomic Atlas of Human Adult Lung Topography"

**Supplementary Fig. 1: H&E stains of snRNA-seq and SNARE-seq2 lung tissue blocks.**

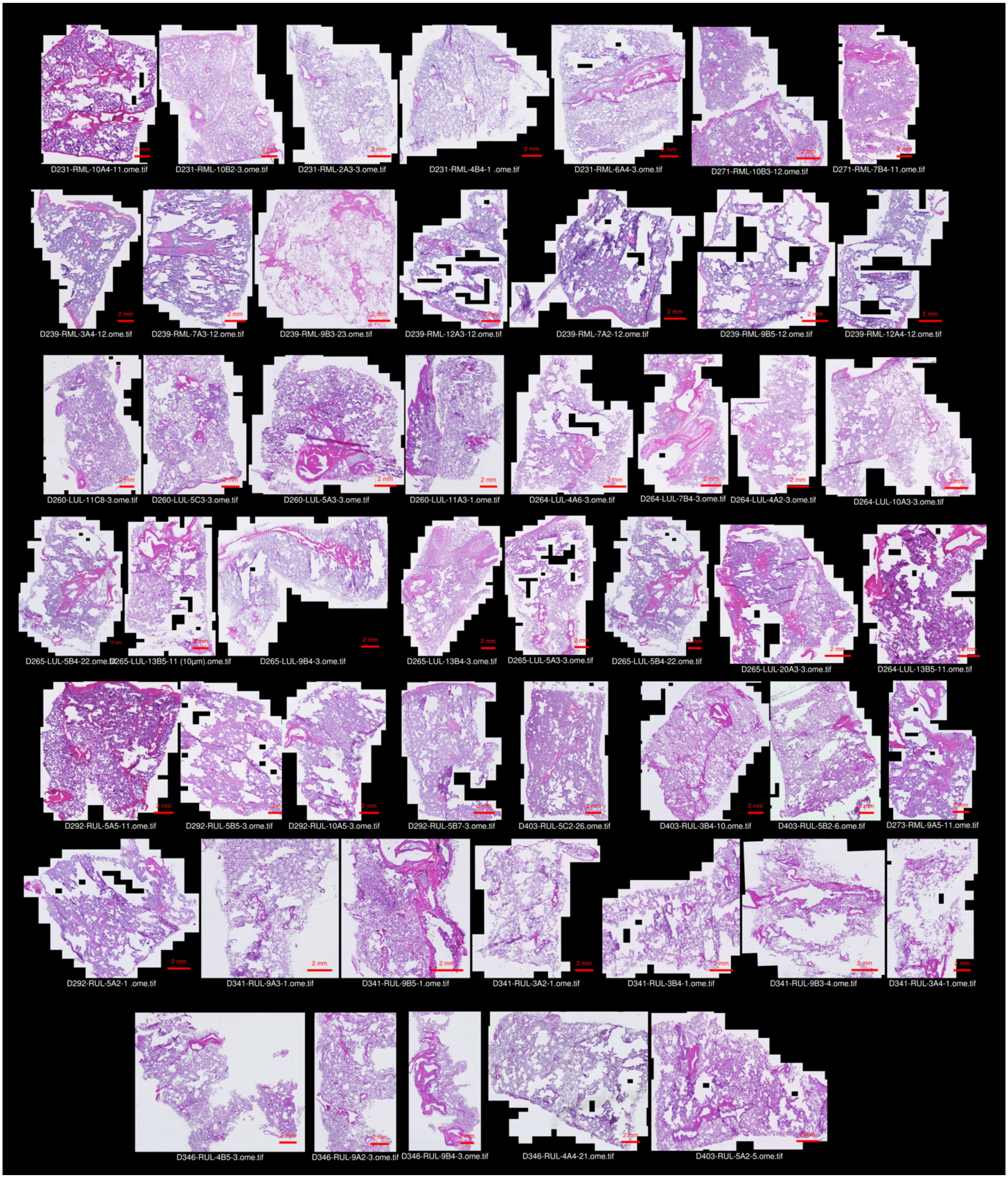


Hematoxylin and eosin stain of frozen, unfixed, optimal cutting media (OCT) embedded tissue sections (10 um thick) serial to the sections (40 um x 8-10) used for snRNA-seq and SNARE2-seq. Scale bars = 2mm.

**Supplementary Fig. 2: H&E stains of FFPE lung tissue blocks.**

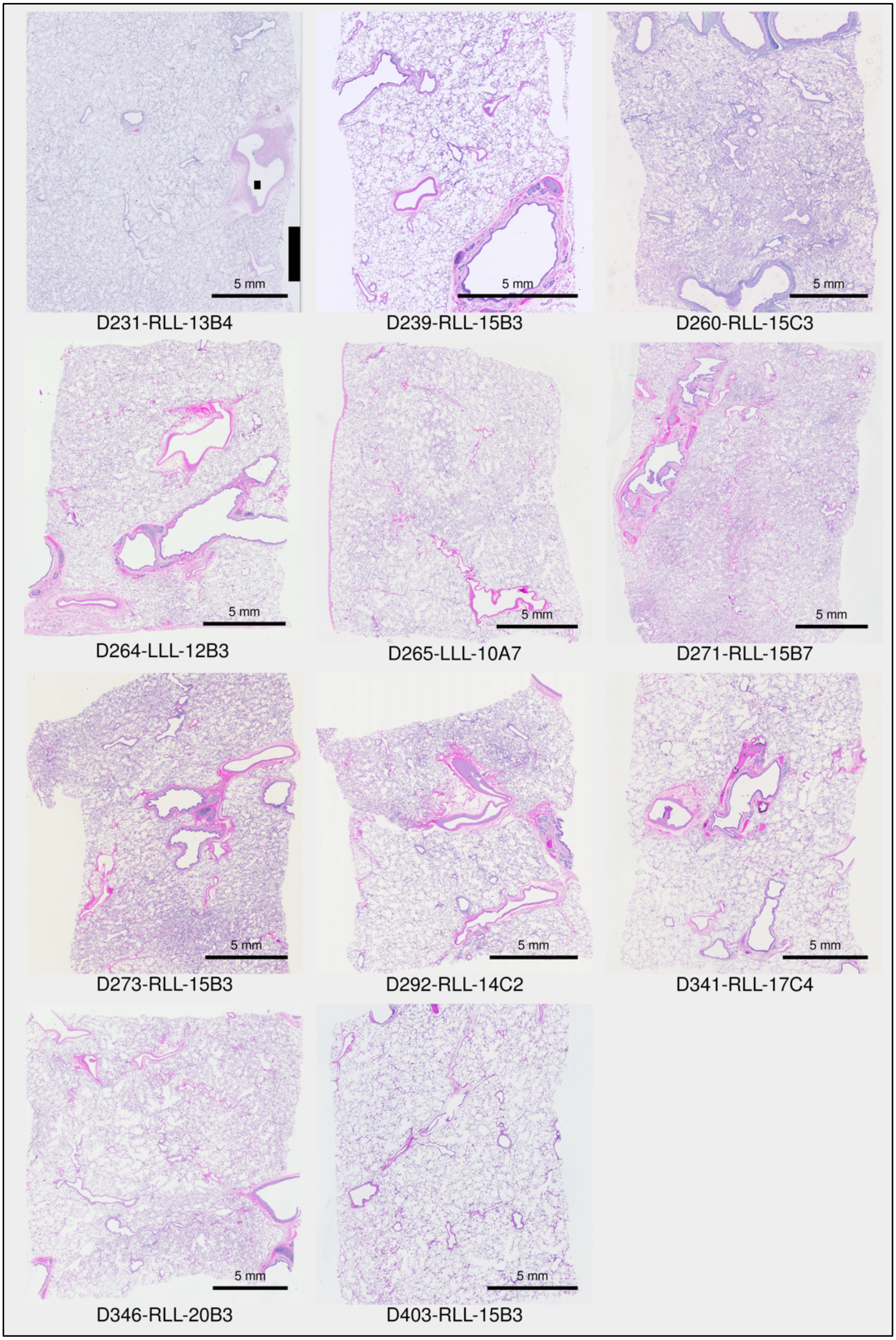

Representative formalin inflated, paraffin embedded (FFPE), H&E sections (5 um) representative of 11 donor lung cases used for single nuclei transcriptomic and open chromatin, spatial transcriptomics and multiplexed immunofluorescence assays. Block IDs are consistent with BRINDL inventory.

**Supplementary Fig. 3**: Spatial proteomics localizes adult pulmonary neuroendocrine cells (PNECs).

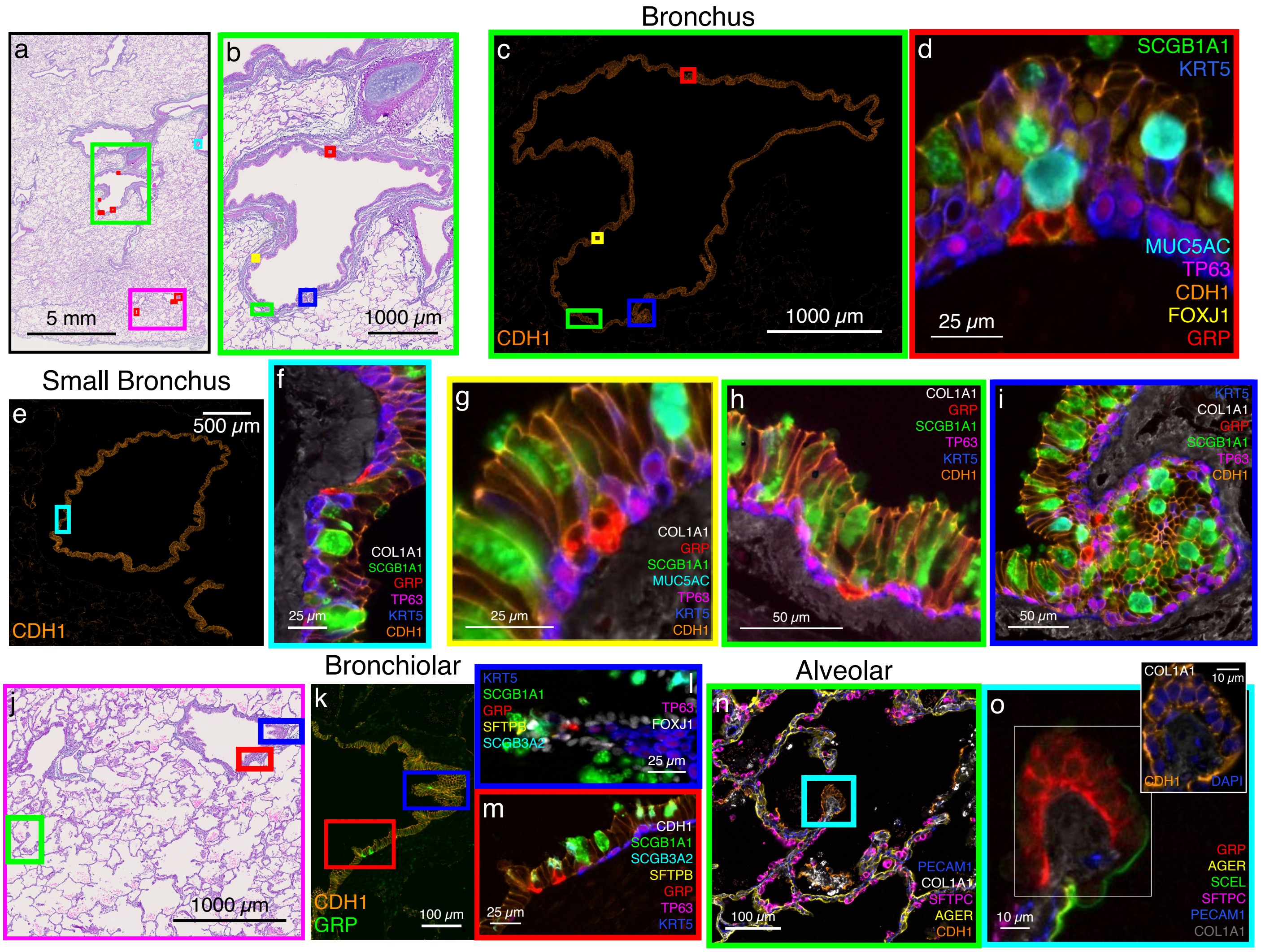


**a**, H&E of whole tissue block, frames correspond to panels **b-c** ,**f**, and **j**; red indicates regions imaged for PNECs**. b**, Digital zoom to panel **a**, green region, frames correspond to panels **d**, and **g-i**. **c** and **e**, Five examples of GRP+ PNECs, in a basal cell location of bronchial airway epithelium expressing MUC5AC+ SCGB1A1+ aw.secretory and TP63+KRT5+ basal cells. **d**, **f**, **g-i**, Digital zoom to frames in **c** and **e**. **j**, Digital zoom to panel **a**, magenta region, frames correspond to panels **k** and **n**. **k-m**, Examples of GRP+ PNECs amongst (**l**)bronchiolar SCGB3A2+SFTPB+ airway region and (**m**) TP63+ KRT5+ basal cells. **n-o**, GRP+ neuroendocrine body collection of PNECs in alveolar region. CDH1/e-caderin (orange) is used throughout to delineate epithelial cell membranes.  Figure is based on D273 and representative of multiple tissue samples.

**Supplementary Fig. 4: MERFISH raw spatial communities identified by BANKSY.**

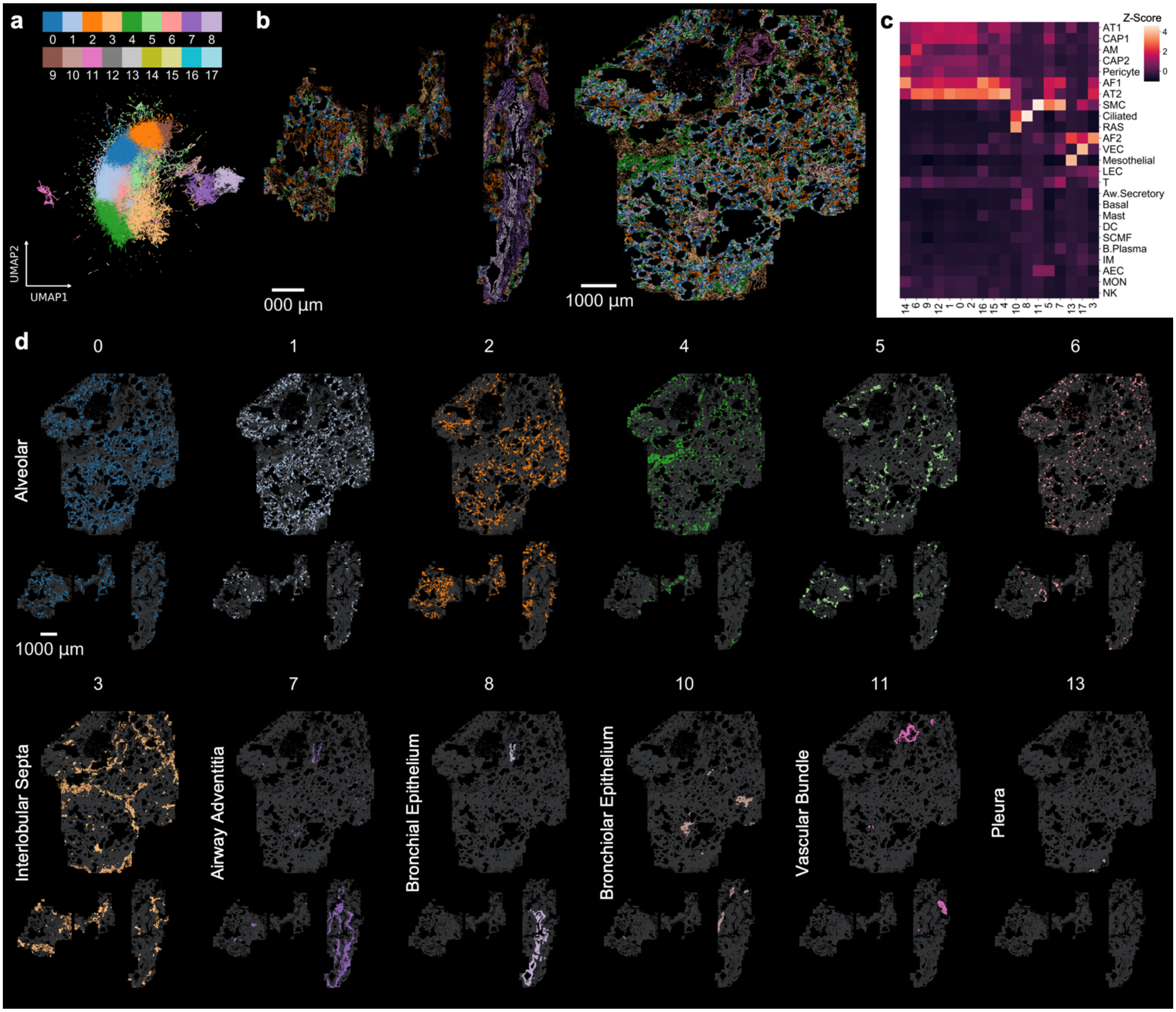


**a**, UMAP projection of MERFISH cells colored by raw BANKSY-labeled clusters (resolution 0.60) before collapsing into named spatial neighborhoods. **b**, Spatial map of cells colored by raw BANKSY-labeled clusters. **c**, Clustered heatmap showing the relationship between cell types and collapsed BANKSY clusters. Values are Z-scored across BANKSY clusters, highlighting relative cell type enrichment within each spatial community and grouping similarly composed BANKSY clusters. **d**, Individual spatial maps of BANKSY clusters by structural neighborhood. The top 6 clusters by cell count are shown for the collapsed Alveolar neighborhood in top row. Clusters 9, 12, 14, 15, 16 not shown. For each of the remaining neighborhoods in the bottom row, except for Interlobular Septa, only one Banksy cluster was assigned. Additional cluster 17 containing only 64 cells was assigned to the Interlobular Septa neighborhood and not shown.
