## Supplementary Tables 1-12 for "A Multimodal Spatial and Epigenomic Atlas of Human Adult Lung Topography": Supplementary Data snHLA.pdf

Supplementary Table Legends

**Supplementary Table 1. Donor Demographics and Clinical/Pathological Assessment.**
Clinical metadata and histopathology for 11 healthy lung tissue donors used in this study.

**Supplementary Table 2. Lung Block Anatomical Characterizations and Use in Assays.**
Summary of omic experiments presented on 63 human lung donor blocks, including location, pathology review, and IDs for HuBMAP Portal data access.

**Supplementary Table 3. snRNA cluster annotations and QC metadata.**
Summary of cell type annotations, number of nuclei, mean UMI, and mean genes per lung block and donor for all snRNA clusters.

**Supplementary Table 4. snHLA Cell Type and Marker Gene Dictionary.**
Cell type annotations at class and different subclass levels with curated and NSForest marker genes for harmonized 10X snRNA and SNARE-RNA.

**Supplementary Table 5. snHLA Cell Type Alignment with Cell Ontology IDs.**

**Supplementary Table 6. snRNA NS-Forest Marker Genes.**
Summary of NS-Forest necessary and sufficient marker and binary genes at each annotation level.

**Supplementary Table 7. Antibodies Used in Multiplexed Immunofluorescence.**
Description of target protein antibodies used in MxIF panels including dilution factors.

**Supplementary Table 8. SNARE2 RNA/AC cluster annotations and QC metadata.**

**Supplementary Table 9. Subclass level 3 cell type specific accessible regions and transcription factor motifs.**
Differentially Accessible Regions (DARs), top ChromVAR transcription factor motifs, and top active (accessible and expressed) ChromVAR transcription factor motifs for subclass.L3 annotations.

**Supplementary Table 10. MERFISH gene panel.**
List of genes and codebook for MERFISH.

**Supplementary Table 11. Summary of Cellchat receptor ligand interactions.**

**Supplementary Table 12. Subclass level 5 cell type specific accessible regions and transcription factor motifs.**
Differentially Accessible Regions (DARs), top ChromVAR transcription factor motifs, and top active (accessible and expressed) ChromVAR transcription factor motifs for subclass.L5 annotations.
