## Supplementary Tables 1-12 for "A Multimodal Spatial and Epigenomic Atlas of Human Adult Lung Topography": Supplementary Table 1 Donor Demographics and Clinical-Pathological Assessment.docx

Clinical metadata and histopathology for 11 healthy lung tissue donors used in this study.

| **HuBMAP ID** | **Donor**  **ID** | **Age**  **(Yrs)** | **Sex** | **Race/ Ethnicity** | **Cause of Death** | **ARDS Status** | **Histopathological Findings** | **Past Medical History** | **Positive Serology** | **Wt (kg)** | **BMI** | **WIT (hrs)** | **CIT (hrs)** |
| --- | --- | --- | --- | --- | --- | --- | --- | --- | --- | --- | --- | --- | --- |
| HBM957. XGCL.285 | D273 | 19.9 | M | White | Drug Intox | neg | Normal lung structure; Pulmonary arterial thrombus; Venous micro-thrombi; Patchy acute inflammation, atelectasis, and edema | Chronic non-IV drug use | CMV IgG | 77 | 21.8 | 0 | 34 |
| HBM849. FBJV.474 | D265 | 23.8 | M | White | Brain Injury | mild | Normal lung structure; Patchy mild peri airway lymphocytic inflammation and macrophage accumulation; Patchy mild fibrin | None | CMV IgG, EBV IgG & IgM | 75 | 21.3 | 0 | 32 |
| HBM626. JCQZ.225 | D260 | 25.0 | M | White | Brain Injury | neg | Normal lung structure; Mild to moderate acute bronchiolitis and organizing hemorrhage | Chronic sinusitis | EBV IgG | 90 | 28.5 | 0 | 33 |
| HBM994. NXZC.854 | D292 | 25.0 | F | White | Drug Intox | neg | Normal alveolar structure; Mild airway smooth muscle thickening; Focal mucostasis; Mild peri airway and perivenular lymphocytic inflammation; Patchy acute hemorrhage | None; Depressant drug use | CMV IgG, EBV IgG | 83 | 30.2 | 0 | 32 |
| HBM852. DHHR.655 | D264 | 25.2 | M | Hispanic | Stroke | neg | Normal lung architecture; Scattered aggregates of pigmented macrophages | None; Daily marijuana | EBV IgG | 70 | 20.9 | 0 | 21 |
| HBM443. VFRD.453 | D231 | 33.0 | F | Black/AA | Anoxic Brain Injury | neg | Normal alveolar growth and structure; Patchy mild peri airway fibrosis, increased smooth muscle; Focal prominence of bronchial goblet cells; Increased alveolar macrophages; Patchy mild mixed inflammation; Mild intimal hyperplasia, large pulmonary artery branches; Scattered microvascular thrombi | Stable, chronic ventilation due to brain injury; Renal calculi | CMV IgG, EBV IgG | 104 | 35.9 | 0 | 29 |
| HBM943. SCQQ.877 | D239 | 37.0 | M | Black/AA | Stroke | mild | Largely normal alveolar structure; Mild bronchiectasis and goblet cell hyperplasia; Foci of bronchiolar metaplasia; Marked occlusive intimal hyperplasia of bronchial vessels; Medial hypertrophy and intimal hyperplasia of large and medium-sized veins; Scattered hemosiderin laden macrophages | Hypertension, obesity; Tox screen + cannabinoids; | none | 151 | 45.0 | 0 | 19 |
| HBM848. BMDK.429 | D403 | 49.6 | F | Asian | Brain Tumor | neg | Normal alveolar structure; Mild peri airway anthracitic pigment; Mild medial pulmonary arterial hypertrophy and intimal hyperplasia; Rare bone marrow emboli and calcified thrombi; Focal proximal airway squamous metaplasia | Seasonal allergies; Recent non-lung tumor without metastasis | CMV IgG, EBV IgG | 63 | 22.4 | 0 | 11 |
| HBM428. GRPJ.489 | D271 | 53.0 | M | White | Stroke | mod | Normal lung structure; Patchy minimal acute and chronic airway inflammation; Patchy mild acute alveolar inflammation with organizing fibrin; Intimal pulmonary vein fibrosis consistent with aging | None | EBV IgG | 90 | 26.9 | 0 | 32 |
| HBM643. KCCK.866 | D341 | 56.8 | F | Hispanic | Stroke | neg | Normal alveolar structure; Medial hypertrophy and focal intimal fibrosis of large pulmonary arteries; Acute and calcified arterial thrombi as well as rare microvascular thrombi; Intimal pulmonary vein fibrosis; Bronchiolar dilation with peri airway anthracitic pigment and focal mild mucostasis | Hypertension, polycystic kidney disease, breast cancer > 5 yrs prior | CMV IgG, EBV IgG | 48 | 19.5 | 0 | 34 |
| HBM932. JNVS.672 | D346 | 59.0 | M | White | Stroke | mild | Normal alveolar and airway structure; Hypertensive vascular changes with medial hypertrophy, intimal fibrosis | Hypertension | EBV IgG | 70 | 22.6 | 0 | 26 |

All negative for SARSCoV2 infection; Serology (hepatitis A, B and C, syphilis, HIV, CMV, EBV) negative except as indicated. BMI is body mass index, kg/m^2^. ARDS is acute respiratory distress syndrome by Berlin definition (PaO2/FiO2 ratio; Ranieri VM, Rubenfeld GD, Thompson BT, et al. Acute respiratory distress syndrome: the Berlin definition. JAMA. 2012;307:2526–33.) Race and/or ethnicity reported by next of kin. WIT and CIT are warm and cold ischemic times.
