## Supplementary Tables 1-12 for "A Multimodal Spatial and Epigenomic Atlas of Human Adult Lung Topography": Supplementary Table 7 MxIF Antibodies .docx

**Supplementary Table 13: Antibodies Used in Multiplexed Immunofluorescence (MxIF)**

Description of target proteins and antibodies used in MxIF panels including dilution factors.

| **UniProt #** | **HGNC_ID** | **Target Protein** | **Vendor** | **Host (clonality)** | **Isotype** | **Clone Id** | **RRID** | **Dilution Factor** |
| --- | --- | --- | --- | --- | --- | --- | --- | --- |
| P12830 | HGNC:1748 | CDH1 | Akoya | Mouse (m) | IgG1 | 4A2C7 | AB_2895057 | 200 |
| P02452 | HGNC:2197 | COL1A1 | Abcam | Mouse (m) | IgG3 | 3G3 | AB_2081873 | 100 |
| P46013 | HGNC:7107 | MKI67 | Akoya | Mouse (m) | IgG1 | B56 | AB_2895046 | 200 |
| P62736 | HGNC:130 | ACTA2 | Akoya | Mouse (m) | IgG2a | 1A4 | AB_2936084 | 200 |
| P11686 | HGNC:10802 | SFTPC | Invitrogen | Rabbit (p) | IgG |  | AB_2717696 | 500 |
| P01730 | HGNC:1678 | CD4 | Akoya | Rabbit (m) | IgG | EPR6855 | AB_2915936 | 200 |
| P34810 | HGNC:1693 | CD68 | Akoya | Mouse (m) | IgG1 | KP1 | AB_2935894 | 200 |
| P08575 | HGNC:9666 | PTPRC | Akoya | Rabbit (m) | IgG | D9M8I | AB_2915946 | 200 |
| P29017 | HGNC:1636 | CD1C | Novus | Mouse (m) | IgG1 | OTI2F4 | AB_3083490 | 50 |
| P07766 | HGNC:1674 | CD3E | Akoya | Rabbit (m) | IgG | EP449E | AB_2936080 | 200 |
| P11836 | HGNC:7315 | MS4A1 | Akoya | Mouse (m) | IgG2a | L26 | AB_2915939 | 200 |
| P16284 | HGNC:8823 | PECAM1 | Akoya | Rabbit (m) | IgG | EP3095 | AB_2915935 | 200 |
| P02776 | HGNC:8861 | PF4 | PeproTech | Rabbit (p) | IgG |  | AB_147868 | 200 |
| Q9Y5Y7 | HGNC:14687 | LYVE1 | R&D Systems | Goat (p) | IgG |  | AB_355144 | 100 |
| P01732 | HGNC:1706 | CD8A | Akoya | Mouse (m) | IgG1 | C8/144B | AB_2915960 | 200 |
| P01903 | HGNC:4947 | HLA-DRA | Akoya | Rabbit (m) | IgG | EPR3692 | AB_2928988 | 200 |
| Q15109 | HGNC:320 | AGER | Abcam | Rabbit (m) | IgG | EPR21171 | AB_2884897 | 100 |
| Q9H3D4 | HGNC:15979 | TP63 | Abcam | Rabbit (m) | IgG | EPR5701 | AB_3083495 | 100 |
| Q15661 | HGNC:12019 | TPSAB1 | Abcam | Mouse (m) | IgG1 | AA1 | AB_303023 | 1000 |
| P08571 | HGNC:1628 | CD14 | Akoya | Rabbit (m) | IgG | EPR3653 | AB_3083457 | 200 |
| P54709 | HGNC:806 | ATP1A1 | Abcam | Rabbit (m) | IgG | EP1845Y | AB_2890241 | 100 |
| Q86VB7 | HGNC:1631 | CD163 | Akoya | Rabbit (m) | IgG | EPR19518 | AB_2935895 | 200 |
| P02462 | HGNC:2202 | COL4A1 | Akoya | Rabbit (m) | IgG | EPR20966 | AB_2927676 | 200 |
| P13647 | HGNC:6442 | KRT5 | Akoya | Rabbit (m) | IgG | EP1601Y | AB_3083458 | 200 |
| Q9BZS1 | HGNC:6106 | FOXP3 | Akoya | Mouse (m) | IgG1 | 236A/E7 | AB_2927679 | 200 |
| P20702 | HGNC:6152 | ITGAX | Akoya | Mouse (m) | IgG2b | 118/A5 | AB_3083459 | 200 |
| P05164 | HGNC:7218 | MPO | Akoya | Rabbit (m) | IgG | E1E7I | AB_2927678 | 200 |
| O95171 | HGNC:10573 | SCEL | Abcepta | Rabbit (p) | IgG |  | AB_10818433 | 100 |
| Q13509 | HGNC:20772 | TUBB3 | R&D | Mouse (m) | IgG2a | TUJ-1 | AB_357520 | 400 |
| P11684 | HGNC:12523 | SCGB1A1 | R&D | Rat (m) | IgG1 | 394324 | AB_2183286 | 400 |
| Q96PL1 | HGNC:18391 | SCGB3A2 | Abcam | Rabbit (m) | IgG | EPR11463 | AB_3083498 | 400 |
| P98088 | HGNC:7515 | MUC5AC | Abcam | Mouse (m) | IgG1 | 45M1 | AB_3083499 | 100 |
| Q92786 | HGNC:9459 | PROX1 | Abcam | Rabbit (m) | IgG | EPR19273 | AB_2894898 | 200 |
| P02533 | HGNC:6416 | KRT14 | Akoya | Rabbit (p) | IgG | Poly19053 | AB_3095339 | 200 |
| Q92949 | HGNC:3816 | FOXJ1 | ThermoFischer | Mouse (m) | IgG1 | 2A5 | AB_1548835 | 500 |
| P07492 | HGNC:4605 | GRP | LSBio | Rabbit (p) | IgG |  | AB_3096183 | 500 |
| P07988 | HGNC:10801 | SFTPB | Invitrogen | Rabbit (p) | IgG |  | AB_2609628 | 50 |
| P08174 | HGNC:2665 | CD55/DAF | R&D Systems | Goat (p) | IgG |  | AB_355106 | 300 |
| Q86YL7 | HGNC:29602 | PDPN | Akoya | Mouse (m) | IgG2a | NC-08 | AB_3082979 | 200 |
| Q03135 | HGNC:1527 | CAV1 | Akoya | Rabbit (m) | IgG | D46G3 | AB_3508120 | 200 |
| Q9HC84 | HGNC:7516 | MUC5B | Abcam | Rabbit (p) | IgG |  | AB_10712492 | 75 |
| P48431 | HGNC:11195 | SOX2 | Akoya | Rabbit (m) | IgG | SP76 | AB_3094504 | 200 |
| P48436 | HGNC:11204 | SOX9 | Cell Signaling | Rabbit (m) | IgG | D8G8H | AB_2665492 | 1000 |
| P05787 | HGNC:6446 | KRT8 | BioLegend | Mouse (m) | IgG2a | 1E8 | AB_2616821 | 400 |
| P04264, P35908, P12035, P19013, P13647, P02538, P04259, P48668, P08729, P05787, P13645, P02533, P19012, P08779, P08727 | HGNC:6412, HGNC:6439, HGNC:6440, HGNC:6441, HGNC:6442, HGNC:6443, HGNC:6444, HGNC:20406, HGNC:6445, HGNC:6446, HGNC:6413, HGNC:6416, HGNC:6421, HGNC:6423, HGNC:6436 | KRT1, KRT2, KRT3, KRT4, KRT5, KRT6A, KRT6B, KRT7, KRT8, KRT10, KRT14, KRT15, KRT16, KRT19 | Akoya | Mouse (m) | IgG1 | AE-1/3 | AB_3083456 | 200 |

RRID at [www.antibodyregistry.org](http://www.antibodyregistry.org); Clonality: (m) monoclonal, (p) polyclonal
